# Feeding Kinematics And Morphology Of The Alligator Gar (*Atractosteus Spatula*, Lacépède, 1803): Feeding Mechanics Of Atractosteus Spatula

**DOI:** 10.1101/561993

**Authors:** Justin B. Lemberg, Neil H. Shubin, Mark W. Westneat

## Abstract

Modern (lepisosteid) gars are a small clade of seven species and two genera that occupy an important position on the actinopterygian phylogenetic tree as members of the Holostei (*Amia* + gars), sister-group of the teleost radiation. Often referred to as “living fossils,” these taxa preserve many plesiomorphic characteristics used to interpret and reconstruct early osteichthyan feeding conditions. Less attention, however, has been paid to the functional implications of gar-specific morphology, thought to be related to an exclusively ram-based, lateral-snapping mode of prey capture. Previous studies of feeding kinematics in gars have focused solely on members of the narrow-snouted *Lepisosteus* genus, and here we expand that dataset to include a member of the broad-snouted sister-genus and largest species of gar, the alligator gar (*Atractosteus spatula*, Lacépède, 1803). High-speed videography reveals that the feeding system of alligator gars is capable of rapid expansion from anterior-to-posterior, precisely timed in a way that appears to counteract the effects of a bow-wave during ram-feeding and generate a unidirectional flow of water through the feeding system. Reconstructed cranial anatomy based on contrast-enhanced micro-CT data show that a lateral-sliding palatoquadrate, flexible intrasuspensorial joint, pivoting interhyal, and retractable pectoral girdle are all responsible for increasing the range of motion and expansive capabilities of the gar cranial linkage system. Muscular reconstructions and manipulation experiments show that, while the sternohyoideus is the primary input to the feeding system (similar to other “basal” actinopterygians), additional input from the hyoid constrictors and hypaxials play an important role in decoupling and modulating between the dual roles of the sternohyoideus: hyoid retraction (jaw opening) and hyoid rotation (pharyngeal expansion) respectively. The data presented here demonstrate an intricate feeding mechanism, capable of precise control with plesiomorphic muscles, that represents one of the many ways the ancestral osteichthyan feeding mechanism has been modified for prey capture.

**RESEARCH HIGHLIGHTS:** Alligator gars use a surprisingly expansive cranial linkage system for prey capture that relies on specialized joints for increased mobility and is capable of precise modulation from anterior to posterior using plesiomorphic osteichthyan musculature.

## 1 INTRODUCTION

Despite the small number of extant gar species, modern (lepisosteid) gars have long been recognized as a scientifically important clade due to their conserved morphology and, more recently, their phylogenetic placement within Holostei (gars + *Amia*), sister-taxon to modern teleosts (Grande, 2010; Near et al., 2012). As such, gars provide an important outgroup comparison in phylogenetic studies for approximately half of vertebrate diversity (Near et al., 2012). Thanks in part to their uniquely low rates of speciation and morphological evolution (Rabosky et al., 2013), gars preserve many plesiomorphic features independently lost in other groups that have proven invaluable for inferring ancestral conditions for osteichthyans (actinopterygians + sarcopterygians) as a whole. Biomechanical studies have used similarities between gars and other non-teleost, non-tetrapod osteichthyans to infer ancestral conditions of the early osteichthyan feeding mechanism (Bemis & Lauder, 1986; Carroll & Wainwright, 2003; Lauder, 1980a, 1980b; Wilga, Wainwright, & Motta, 2000), while developmental studies have used their conserved gene sequences to identify homologous regions of genetic code between teleosts and tetrapods (Braasch et al., 2016; Nakamura, Gehrke, Lemberg, Szymaszek, & Shubin, 2016). Often described as “living fossils” (Darwin, 1859; Grande, 2010; Rabosky et al., 2013; Wiley, 1976; Wright, David, & Near, 2012), gars have helped to bridge our understanding of the evolution of modern forms to the distant past.

However, while their plesiomorphic trait retention is useful, it is important to note that since diverging from the rest of Neopterygii approximately 300 million years ago (Near et al., 2012) gars have accumulated numerous specialized features, the functional implications of which are not fully understood. Some features relate to gars being the last remaining holdovers of an ancient, adaptive radiation, the ginglymodians (Cavin, 2010; Grande, 2010; Schaeffer, 1967), whereas others presumably relate to their unique method of capturing prey using rapid lateral strikes (Lauder & Norton, 1980; Muller, 1989; Porter & Motta, 2004). These specializations include platyrostral crania (i.e., flat-snouted), elongate jaws, anteriorly positioned jaw joints, numerous plicidentine teeth, teeth-bearing lacrimomaxillary bones, enlarged palates, broad basipterygoid-metapterygoid processes, dorsally-exposed ectopterygoids, and fusiform bodies with posteriorly-placed midline fins (Arratia & Schultze, 1991; Grande, 2010; Jollie, 1984; Kammerer, Grande, & Westneat, 2006; Lauder, 1980a; Porter & Motta, 2004; Webb, Hardy, & Mehl, 1992; Wiley, 1976). Gars are typically characterized as representing morphological “extremes” towards biting-based, ram-feeding prey capture, but they are consistently outliers in terms of kinematics in comparison with other piscivorous ambush predators with similar body-shapes, such as barracuda, needlefish, and pike (Muller, 1989; Porter & Motta, 2004; Webb et al., 1992). In order to help understand the functional implications of some of the more unusual features of gar anatomy, it is necessary to expand the current dataset of comparative feeding kinematics to a broader taxonomic range of extant gar species.

To further expand our understanding of feeding kinematics within lepisosteids, this study focuses on a species of gar with unique morphology and ecology, the alligator gar, *Atractosteus spatula*. With the most brevirostral jaws (i.e., blunt-snouted) and largest adult body size (Grande, 2010; Kammerer et al., 2006), *A. spatula* differs morphologically from other species of extant gars for which feeding kinematics are available – *Lepisosteus oculatus* (Lauder, 1980a; Lauder & Norton, 1980), *L. platyrhinchus* (Porter & Motta, 2004), and *L. osseus* (Webb et al., 1992). These morphological differences coincide with a generalist diet reported to include fish, smaller gars, invertebrates, birds, and opportunistically scavenged detritus (Grande, 2010; Kammerer et al., 2006; Raney, 1942; Robertson, Zeug, & Winemiller, 2008). Although rostral proportions and diet are known to vary between sympatric species of gars (Grande, 2010; Kammerer et al., 2006; Robertson et al., 2008; Wright et al., 2012), it is uncertain how differing cranial morphology affects the overall patterns of gar feeding kinematics. Therefore, the first aim of this study is to quantify feeding kinematics in *A. spatula*, using high-resolution, high-speed videography, to assess if broad, flat jaws have an effect on the prey strike kinematics and feeding strategies of alligator gars.

It is possible that variation in feeding morphology relates to differences in prey capture methods between gar species, but without an effective mechanism for suction-based prey capture it is hard to hypothesize alternate feeding strategies. Although gars are known to use suction for prey-processing, (Lauder & Norton, 1980; Werth, 2006), suction-generation for the purposes of prey capture is thought to be minimal or non-existent due to limitations of the expansive capabilities of the gar feeding apparatus (Lauder, 1980a; Muller, 1989; Porter & Motta, 2004). The elongate jaws of gars with unoccluded gape and fixed premaxillae are unconducive for effectively directing suction-feeding forces during the prey strike (Lauder, 1980a; Wainwright, McGee, Longo, & Patricia Hernandez, 2015). Furthermore, the short hyoid bars and platyrostral morphology of the upper jaws in gars appears to limit suspensorial abduction, one of the key components of volumetric expansion in osteichthyans (Lauder, 1980a; Liem, 1978; Muller, 1989). While gars possess several derived features that could function during suspensorial abduction, such as a patent ectopterygoid-dermatocranium sulcus and a unique sliding metapterygoid-basipterygoid articulation (Arratia & Schultze, 1991; Grande, 2010; Jollie, 1984; Wiley, 1976), with a tight anterior articulation of the palate, laterally positioned jaw joints, and horizontal long axis of the suspensorium (Adriaens & Verraes, 1994; Alexander, 1970; Grande, 2010; Lauder, 1980a), the mechanism for cranial expansion in gars is not fully understood. Therefore, a secondary aim of this study is to document possible mechanisms of cranial expansion in *Atractosteus spatula*, using detailed anatomical reconstructions of dermal and endochondral cranial anatomy based on contrast-enhanced, X-ray micro computed tomography (μCT).

Although gars are thought to be limited in mechanisms of suction-generation, previous feeding kinematics studies have noted gars maintain an anterior-to-posterior expansion of the buccopharyngeal cavity during the prey strike, particularly a pronounced delay in hyoid depression relative to jaw opening (Lauder, 1980a; Lauder & Norton, 1980; Porter & Motta, 2004). While delayed hyoid depression is not unusual in aquatic vertebrates with multiple mechanical pathways for jaw opening, gars are only known to possess one – i.e., the plesiomorphic, osteichthyan jaw opening mechanism, which is a mechanical coupling between the hyoid arch and mandible whereby posterodorsal forces imparted to the hyoid by the sternohyoideus are transmitted to the lower jaw via the mandibulohyoid ligament (Lauder, 1980a; Lauder & Shaffer, 1993; Wilga et al., 2000). Gars lack the specialized jaw opening musculature of derived osteichthyan groups, such as the levator operculi found in *Amia* and teleosts or the depressor mandibulae found in lepidosirenid lungfish or tetrapods (Bemis, 1987; Lauder, 1980a; Lauder & Shaffer, 1993), and, furthermore, gars lost the plesiomorphic coracomandibularis coupling frequently used as either a primary or secondary mechanism of jaw opening in many gnathostome groups (Wilga et al., 2000). Therefore, movement of the hyoid and lower jaws in this system are mechanically linked, and with no other known muscular contributions attaching to the lower jaws in gars (Lauder, 1980a, 1982; Lauder & Norton, 1980), the separate movements of the hyoid apparatus and lower jaws in gars are not fully understood. A third aim of this study is to better understand the jaw opening mechanism of *Atractosteus spatula* by using high-speed videography, contrast-enhanced μCT, and *in situ* manipulations of fresh specimens to help clarify how the feeding mechanism of osteichthyans operates with a plesiomorphic complement of muscles.

## 2 MATERIALS AND METHODS

### Fish specimens and care

*Atractosteus spatula* specimens were obtained from the U.S. Fish and Wildlife Service Warm Springs National Fish Hatchery (Trip Number: 14-023), with all aquarium maintenance and care performed at The University of Chicago under Institutional Animal Care and Use Committee protocol 72365. The fish were trained to feed on pieces of freeze dried krill held in forceps in the presence of bright lights to acclimate fish for later high-speed videography. Although gars are primarily piscivorous, invertebrates form a portion of the diet young gars in the wild (Buckmeier, 2008). The gars were enticed to feed in midwater to enhance their visibility during feeding. Specimens were fed three times a week, with a fasting period of 48-72 hours observed before filming.

### High speed videography and digitizing

A cohort of five actively feeding *A. spatula* juveniles (1.5 years old) were selected for high-speed videography. A mirror submerged in the tank and angled at 45° provided simultaneous lateral and dorsal views. Strike videos were recorded at 500 frames per second using a Photron FASTCAM SA7 color high-speed video camera with a shutter speed of 1/1000^th^ of a second. A total of 101 feeding strikes was recorded. Strikes were screened for clarity and positioning of the fish relative to both views of the camera, and 25 of the best videos were selected for digitization, with five videos per individual forming the basis of the kinematics dataset.

Videos were digitized using the StereoMorph R package version 1.5.1 (Olsen & Westneat, 2015). Seven landmarks in dorsal view and eleven landmarks in lateral view were digitized in each frame from the onset of the feeding strike to the completion of the reset phase (Fig. 1). In instances when a landmark was occluded (e.g., by the forceps, prey item, or moving out of the field of view), the landmark’s position was interpolated from the position of surrounding markers or discarded until it could be tracked again.

**Figure 1.**
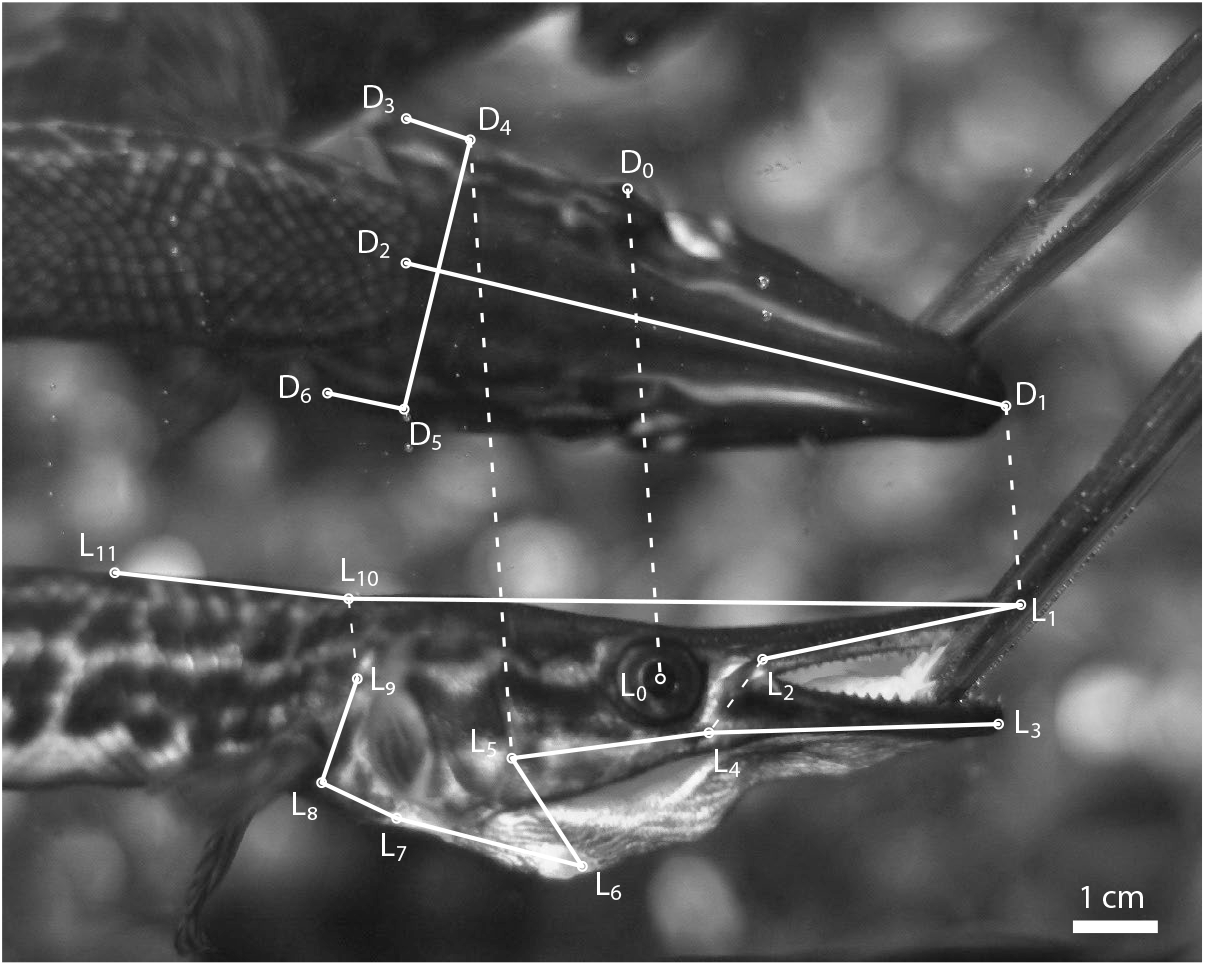
Digitizing schema of anatomical landmarks for high-speed feeding videos. Seven dorsal and eleven lateral points were tracked continuously in StereoMorph throughout the feeding sequence. Dorsal points: center of orbit (D^0^), tip of the upper jaws (D^1^), back of the skull (D^2^), right opercular flap (D^3^), right subopercle-preopercle joint (D^4^), left subopercle-preopercle joint (D^5^), left opercular flap (D^6^) (note the inversion of left and right because the dorsal view is a mirrored view). Lateral points: center of the orbit (L^0^), tip of upper jaws (L^1^), base of upper jaws (L^2^), tip of lower jaws (L^3^), jaw joint (L^4^), right subopercle-preopercle joint (L^5^), anterior tip of the hyoid (L^6^), anterior tip of cleithrum (L^7^), base of right pectoral fin (L^8^), cleithral-supracleithral joint (L^9^), inflection point of neck (L^10^), fixed dorsal body point (L^11^). Other points such as the starting position of the food item in dorsal and lateral views were measured once to establish prestrike starting conditions (Fig. 2).

The known width and inter-tine spacing (0.083 cm = 1/12^th^ cm) of the forceps provided a reliable scale in each feeding video. The number of pixels per centimeter was measured in the clearest frame of each video, where the forceps were held perpendicular in the field of view. An arbitrary point in the center of the prey item was used at the beginning of the prey strike to establish the starting position of the prey.

#### Measuring kinematic variables

Landmarks first were rotated into a global coordinate system. The center of the eye facing the camera was used as a marker for the global vertical axis by measuring its position in dorsal and lateral views throughout the sequence (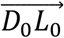, Fig. 1). The angle between this vector 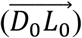 and the global vertical axis was averaged throughout the feeding sequence, and then all landmarks were rotated by that averaged global offset. This aligned all landmarks shared between the dorsal and lateral views in the global vertical axis (Fig. 1) and created a horizontal global axis that is perpendicular to the reflection of the mirror between dorsal and lateral views.

After establishing a global coordinate system, yaw angle was measured and calculated as the offset angle between the midline of the skull in dorsal view 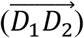 and the global horizontal axis. Yaw greater or less than 0° in dorsal view foreshortens angles in the lateral view by the cosine of its angle (e.g. cos(15°) = 96.59%), so a correction factor, (1/cos(yaw°)), was applied to horizontal distance measurements in lateral view to account for this distortion.

Gape angle was measured as the angle between the upper tooth row 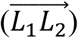, and the tip of the lower jaws to the jaw joint 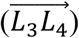 in lateral view. Peak gape was set as time zero in the feeding strike, as it was often associated with other kinematic variables, such as onset of suspensorial abduction and peak lateral-snapping speed (Table 1, Fig. 3).

**Table 1.** Individual and aggregate means of *Atractosteus spatula* feeding kinematics. Means of kinematic variables were calculated for all five individuals with standard deviations (±). ^†^Timing is relative to peak gape at 0 ms. ^‡^Peak value is measured in degrees, except for suspensorial abduction (percent lateral expansion) and snapping/lunging (cm/s). ^§^For hyoid depression, pectoral girdle retraction, and suspensorial abduction, duration only includes time from onset to peak, because the prey is held in the pharyngeal cavity for an extended period of time before prey processing. Sample size: individuals (n = 5), strikes per individual (n = 5).

| Kinematic Variable | Onset (ms) <sup>†</sup> | Peak (ms) <sup>†</sup> | Value at Peak <sup>‡</sup> | Duration (ms) |
| --- | --- | --- | --- | --- |
| Jaw opening/closing | $-17.4 \pm 0.7$ | NA | $48.2 \pm 1.3^\circ$ | $40.7 \pm 3.3$ |
| Neurocranial elevation | $-18.7 \pm 2.1$ | $5.4 \pm 4.8$ | $10.2 \pm 2.2^\circ$ | $53.2 \pm 6.9$ |
| Hyoid depression | $-8.3 \pm 2.8$ | $17.8 \pm 3.6$ | $47.4 \pm 6.0^\circ$ | $26.2 \pm 5.1$ <sup>§</sup> |
| Pectoral girdle retraction | $-6.8 \pm 2.9$ | $26.1 \pm 2.7$ | $19.7 \pm 5.0^\circ$ | $32.9 \pm 2.4$ <sup>§</sup> |
| Suspensorial abduction | $1.0 \pm 2.9$ | $22.4 \pm 2.9$ | $44.9 \pm 7.6\%$ | $21.5 \pm 2.5$ <sup>§</sup> |
| Lateral snapping | $-13.5 \pm 1.5$ | $-2.8 \pm 1.2$ | $143.6 \pm 22.1$ cm/s | $24.4 \pm 0.8$ |
| Forward lunging | $-5.7 \pm 2.2$ | $10.2 \pm 1.7$ | $66.1 \pm 14.1$ cm/s | $118.4 \pm 15.8$ |

**Figure 3.**
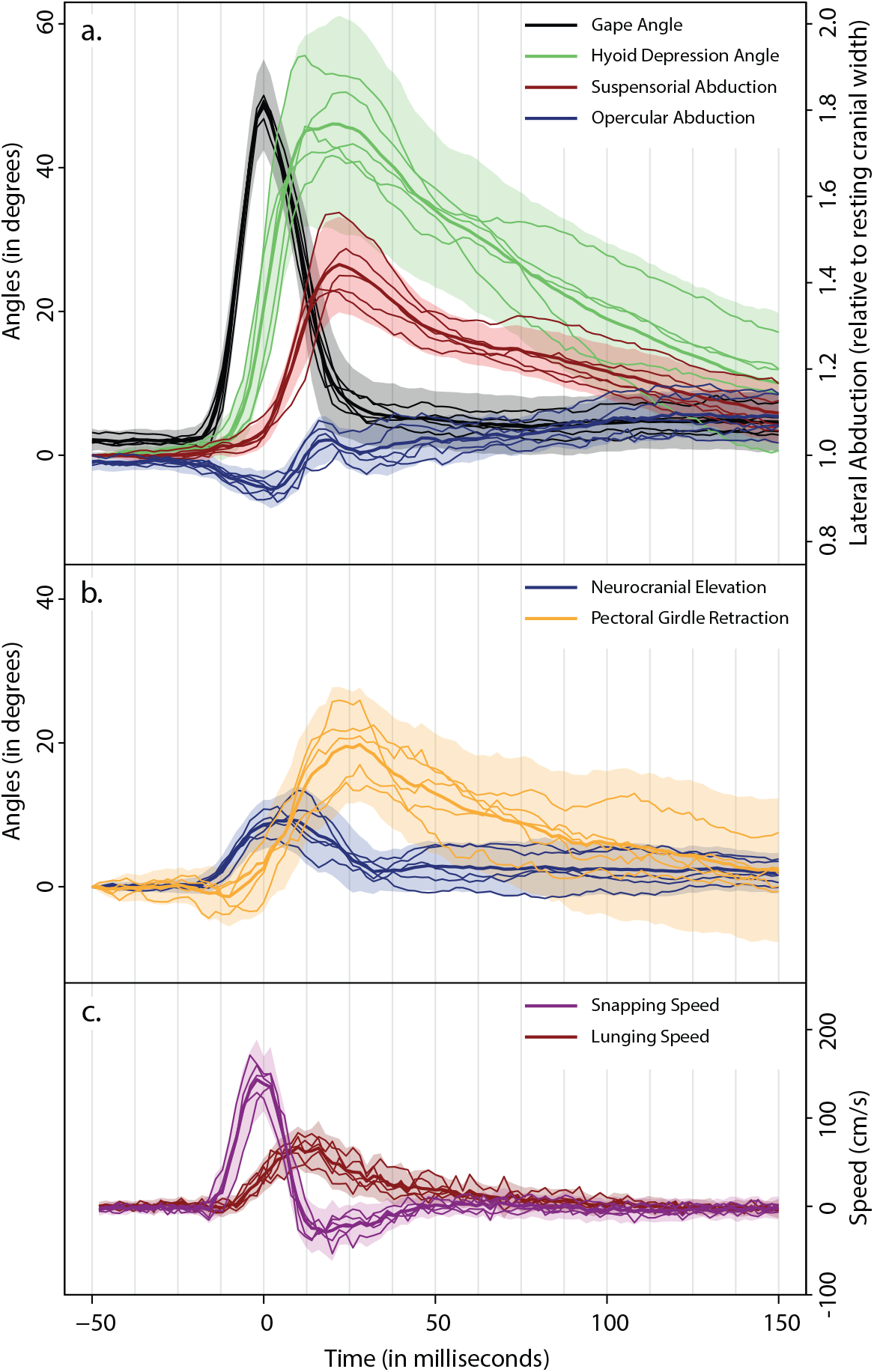
Averaged feeding kinematics of *Atractosteus spatula*. Graphs show the means per individual (thin lines) as well as the aggregate mean (all individuals, thickened line) of each measured kinematic variable. Standard deviations are indicated by the shaded area above and below each aggregate mean. All sequences are aligned with peak gape coinciding with time 0 ms. (*a*) Kinematics show a clear anterior-to-posterior sequence of expansion of cranial elements, with onset of hyoid depression highly variable between individuals (as shown by separation of individual lines for that variable from −10 to 10 ms). (*b*) Neurocranial elevation appears to coincide with the timing of gape angle, whereas pectoral girdle retraction appears closely associated with hyoid depression. (*c*) Lateral-snapping speed promptly decelerates with the onset of hyoid depression. (N.B., kinematics for suspensorial abduction, opercular abduction, snapping speed, and lunging speed are aligned with the right axis).

Hyoid depression was measured as the change in angle (relative to the beginning of the feeding strike) between the jaw joint, the bottom of the opercular joint, and the bulge indicating the hypohyals (∠*L*^4^*L*^5^*L*^6^, Fig. 1). The bottom of the opercular joint (marked by the intersection of the subopercle, preopercle, and adjacent suborbital), while not the exact location of the interhyal-ceratohyal joint, was a consistently clear landmark in the feeding videos and an effective proxy for its location between individuals.

Neurocranial elevation was measured as the change in angle (relative to the beginning of the feeding strike) between the inclination of the skull 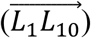 and the body axis 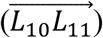. Similarly, pectoral girdle retraction was measured as the change in angle (relative to the beginning of the feeding strike) between the cleithrum 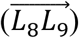 and body axis 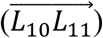.

Suspensorial abduction was measured as the width of the skull at the preopercle-opercle hinge 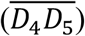 divided by the resting width of the skull in dorsal view. Initially, the preopercle-opercle hinge is not visible in dorsal view, so the widest point of the skull is measured at the sphenotics for the initial values of skull width. This produces a value between 1.0 and 1.7, equal to 0% and 70% expansion of cranial width due to suspensorial abduction, respectively.

Opercular abduction was measured in the same way as suspensorial abduction except, in order to isolate the component of cranial expansion due to opercular abduction alone, the concurrent width of the preopercle-opercle hinge 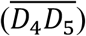 was subtracted from total interopercle width 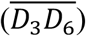. This value was then divided by the resting width of the skull, producing values between 0.82 (opercular adduction) and 1.23 (opercular abduction), equal to 18% constriction and 23% expansion of resting cranial width respectively due to opercular movements independent of suspensorial movements.

Lateral-snapping speed was calculated by first measuring the perpendicular distance of the midline axis of the skull in dorsal view 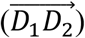 to the initial coordinates of a point in the center of the prey item at the beginning of the prey strike (marked by a ‘+’ in Fig. 2). The distance traversed frame-by-frame in centimeters was multiplied by 500 to get lateral-snapping speed 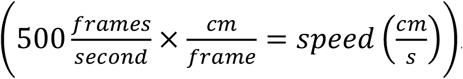. Note that negative values for speed indicate that the center line of the jaws had passed the starting coordinates of the prey.

**Figure 2.**
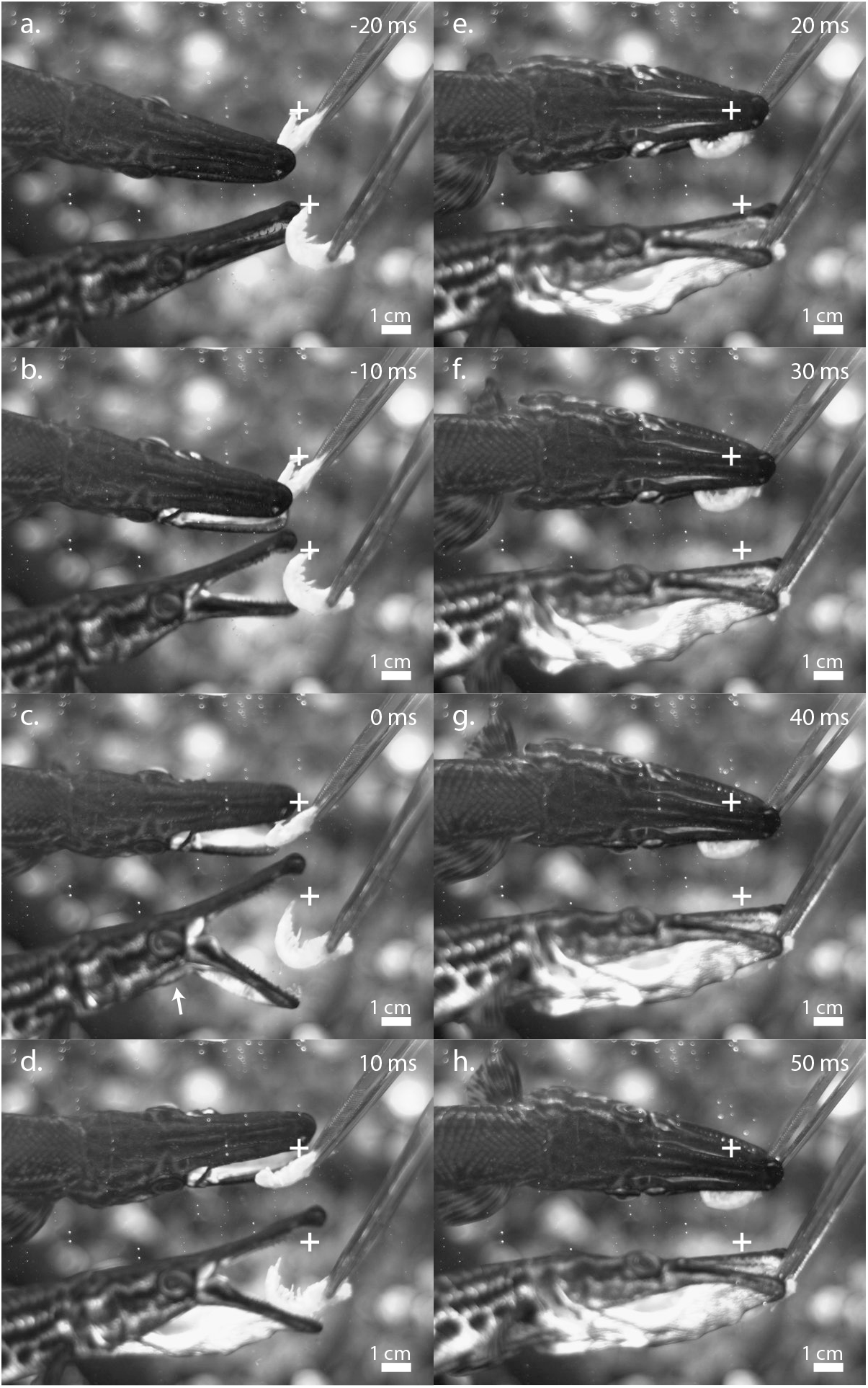
Representative feeding sequence of *Atractosteus spatula* showing suction and delayed hyoid depression. Shown are key frames from high-speed video indicating: (*a*) positioning of the animal prior to strike; (*b*) jaw opening; (*c*) peak gape; (*d*) hyoid depression; (*e*) suspensorial abduction; (*f*) peak pectoral girdle retraction; (*g-h*) kinematic recovery while the opercular flaps are held abducted. The initial starting position of the prey item is marked by a white ‘+’ in both mirror and lateral views. The prey item is drawn into the buccal cavity (*b-c*) prior to the onset of hyoid depression at peak gape (*c*, white arrow), indicating flat-plate suction. Depression of the hyoid, accompanied by retraction of the pectoral girdle, draws the prey item further into the pharyngeal cavity (*d*) just prior to jaw closure (*e*). The scarf joint between the ectopterygoid and frontal bones can be seen as a distinct, white line that appears along the length of the rostrum anterior to the orbits (*e-h*). Peak hyoid depression and suspensorial abduction occurs at 24 ms (not shown).

Forward-lunging speed was measured the same way as lateral-snapping speed, except only the distance between the back of the skull in dorsal view (*D*_2_) and the same initial prey item point was used. The distance traversed frame-by-frame in centimeters was multiplied by 500 to get forward-lunging speed (see above).

### In situ manipulation and preparation for imaging

Individual “D” from the feeding experiments was sacrificed for *in situ* manipulation experiments and detailed anatomical reconstructions based on contrast-enhanced, X-ray micro computed tomography (μCT). The specimen was anesthetized in a solution of tricaine methanesulfonate (MS222), 200 mgL^-1^ buffered with 200 mg sodium bicarbonate (NaHCO^3^), and then euthanized according to procedures approved by the University of Chicago Institutional Animal Care and Use Committee.

*In situ* manipulations were performed to test the full range of cranial kinesis in a fresh specimen. A stainless-steel wire was threaded through the connective tissue directly anterior to the pectoral girdle, allowing the pectoral girdle to be retracted and rotated, which simulated the input of sternohyoideus and hypaxial muscle forces. In addition to these primary inputs to the feeding mechanism, two additional muscle groups were simulated: the hyoid constrictors and jaw adductors. Input of the hyoid constrictor musculature was simulated by holding the hyoid in an elevated position and holding the preopercles in an adducted position while the sternohyoideus and pectoral girdle were retracted. Input of the jaw adductors were simulated by holding the jaws shut while the sternohyoideus and pectoral girdles were retracted. Results were photographed (Fig. 4) and a video recording of this experiment is available online (Supplementary Video 1).

**Figure 4.**
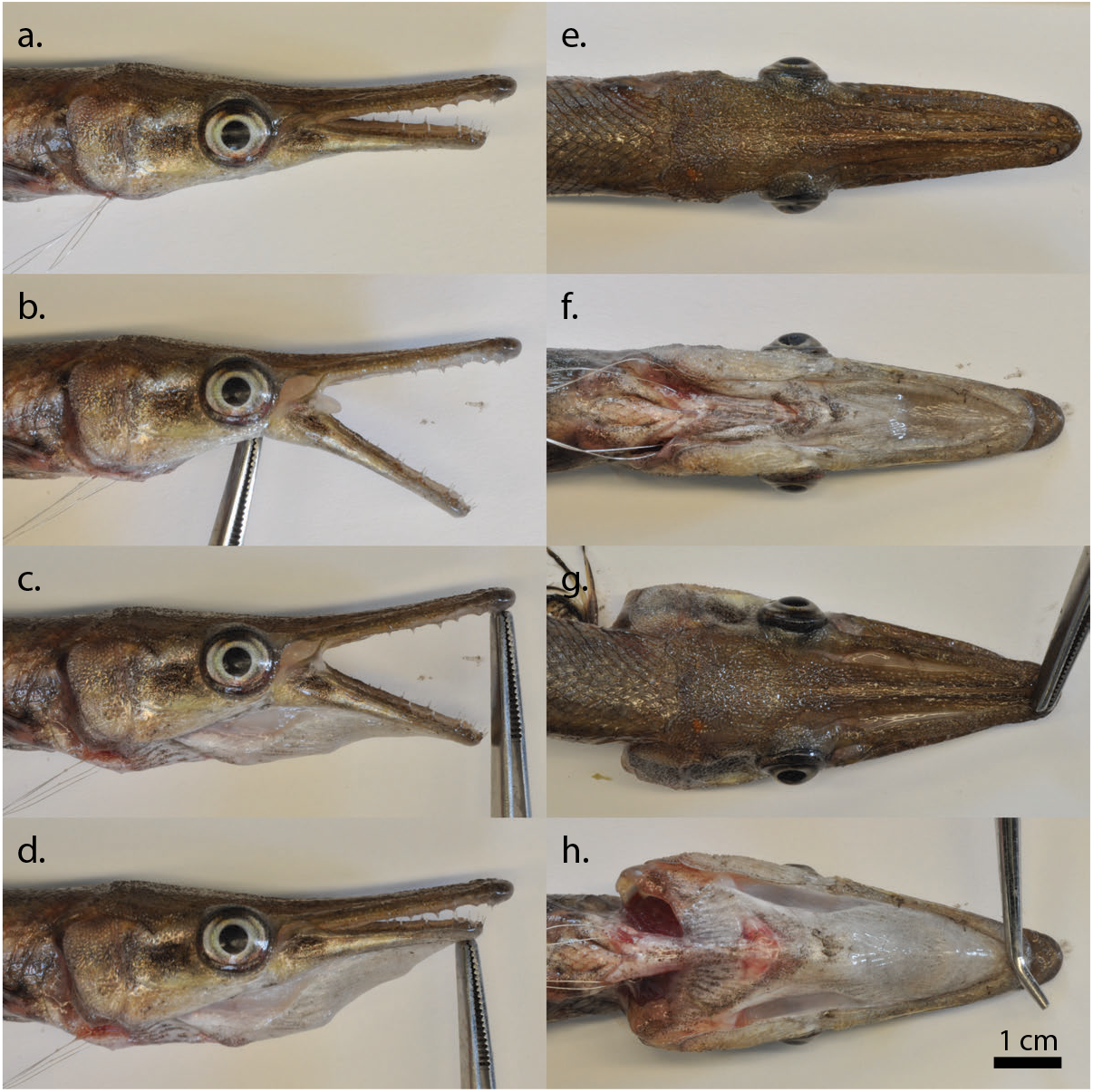
*In situ* manipulation of the *Atractosteus spatula* cranial linkage mechanism. A steel wire looped around the anterior tip of the cleithrum and dorsal to the sternohyoideus is used to simulate input of the sternohyoideus and hypaxial muscles in the feeding system of *A. spatula*. (*a*) At resting state the wire is untensed. (*b*) Tensing the wire with the hyoid held in an elevated state effects jaw opening. (*c*) Tensing the wire with the hyoid unconstrained effects hyoid depression, pectoral girdle retraction, and weak jaw opening. (*d*) Tensing the wire with the jaws held closed effects hyoid depression, pectoral girdle retraction, and flaring of the cheeks (see *g* and *h*). (*e*-*f*) Dorsal and ventral view with wire untensed at resting state. (*g-h*) Dorsal and ventral view with wire tensed and jaws held closed, simulating hyoid rotation and suspensorial abduction. Lateral images (*a-d*) are of the left side reflected to match the orientation of the feeding videos.

After manipulation experiments, the gar was fixed in a solution of 10% formalin in water (weight/volume). The specimen was fixed with fully abducted cheeks and depressed hyoid by filling the pharyngeal cavity with folded paper towel that could easily be removed after the fixation process. No additional method was needed to hold the jaws closed at peak hyoid depression, suggesting the mandibulohyoid ligament is not tensed due to hyoid rotation alone. A period of 1 week in 10% formalin was followed by washing and soaking the specimen for an additional 7 days in water with the pharyngeal cavity empty. Prior to scanning, the specimen was rehydrated in a 20% solution of sucrose in water (weight/volume) for 48 hours.

### Contrast-enhanced μct and segmentation

A reference scan established the position of bones in the specimen. The reference scan was collected using the UChicago PaleoCT (GE Phoenix v|tome|x 240 kv/180 kv scanner) (http://luo-lab.uchicago.edu/paleoCT.html) with the following scanning parameters: voltage = 70 kV, current = 150 μA, timing = 333 ms, voxel width = 25 μm, resolution = 40 pixels/mm, no filter.

Next, the specimen was immersed in a 2.5% solution of phosphomolybdic acid (PMA) in water (weight/volume) for a period of 12 days. During this time, the specimen and solution were set to mix on an orbital shaker, kept at room temperature, and covered with foil to prevent photoreaction. PMA is a contrast agent used to enhance μCT scans (Descamps et al., 2014; Pauwels, Van Loo, Cornillie, Brabant, & Van Hoorebeke, 2013) allowing distinction between a variety of tissues including muscle, bone, ligament, and nervous tissue. The PMA-enhanced μCT scan used the following scanning parameters: voltage = 70 kV, current = 300 μA, timing = 200 ms, voxel width = 25 μm, resolution = 40 pixels/mm, no filter. Some shrinkage occurred between the reference scan and the PMA-enhanced scan, which could pose a problem for measuring muscle volumes, but anatomical structures were readily distinguishable. A preliminary scan after only five days in the PMA solution indicated that PMA had penetrated everywhere except the densest portions of the axial musculature and neural cavity. Twelve days in the PMA solution was sufficient time for the PMA to diffuse throughout the entire body.

#### Digital segmentation

*Atractosteus spatula* μCT data were segmented using Amira 6.2.0 (FEI). To facilitate segmentation, an 8-bit, down-sampled version of the dataset was used to make the initial segmentation at a resolution of 20 pixels/mm. The higher resolution (40 pixels/mm, 16-bit) dataset and reference scan were used to confirm any ambiguous segmentation made in the lower resolution scan.

The digital dissection emphasized bones, muscles, and cartilage of interest for elucidating the feeding mechanism. Visualization incorporated volume renderings of the individually segmented elements, with color-coding based on the region of the braincase the element belonged to: muscles were given a red-to-yellow-to-white color scheme, dermal bones were colored grey-to-white, while endochondral bones and closely associated dermal bones (such as the lacrimomaxillas, and dermal bones of the lower jaws) were given a blue-to-cyan-to-grey-to-white color scheme. Ligaments were left white. Application of these color transitions was made to emphasize the anatomical structures of cranial elements, not strictly to show the distinction between bone and cartilage. Abbreviations used in anatomical figures are listed in Table 2.

**Table 2.**
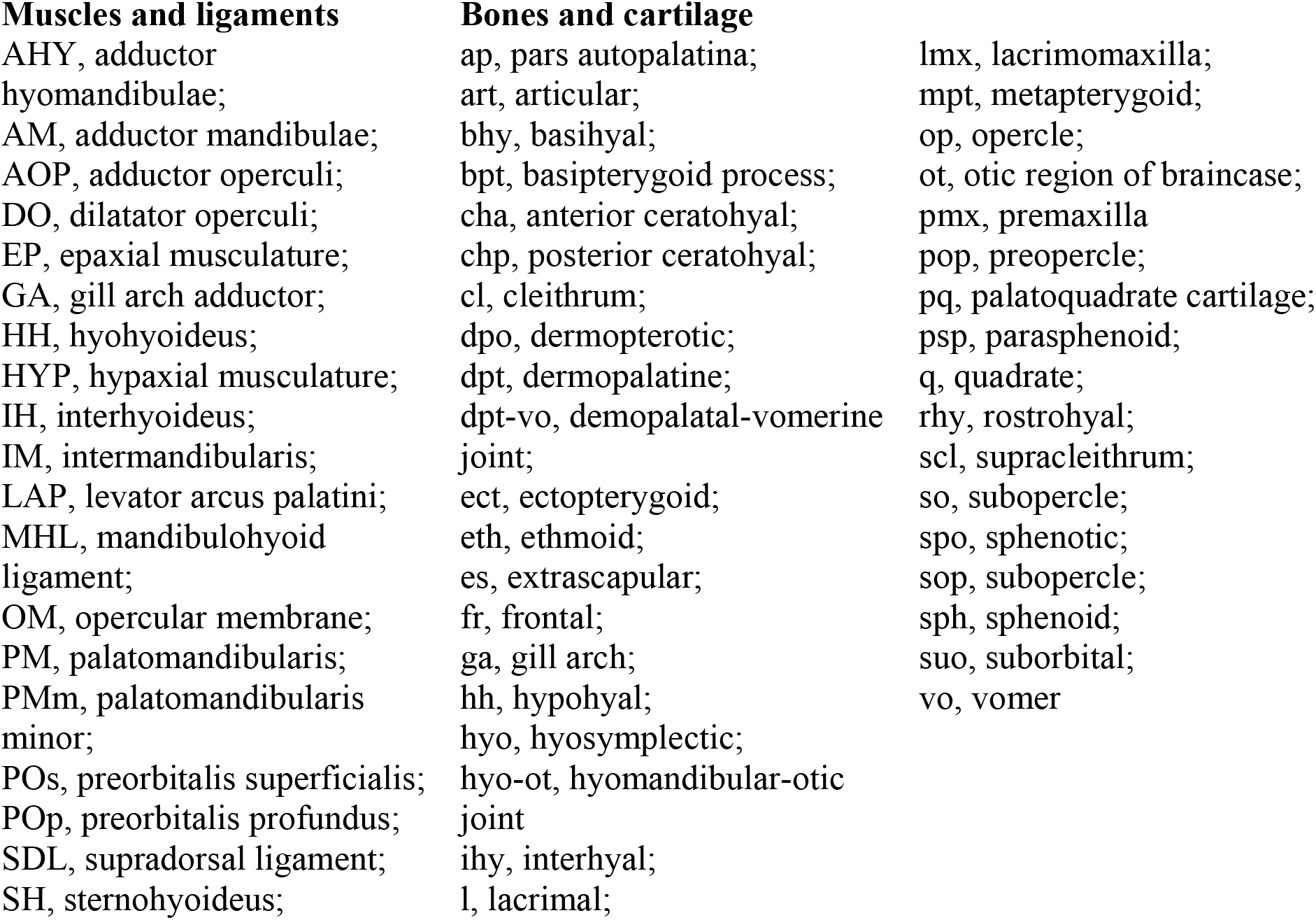
Anatomical abbreviations for *Atractosteus spatula* figures

### Animating the kinematic model

In order to recreate the cranial movements seen in the high-speed videos, individually segmented elements of the cranial skeleton were rotated in Amira 6.2.0 (FEI) into four successive kinematic states: resting-state, jaw opening prior to hyoid depression, hyoid depression prior to jaw closing, and peak suspensorial abduction/hyoid depression. In Amira, each segmented element can be rotated relative to surrounding cranial elements, and the resulting rotation is exported as a (4 × 4) transformation matrix which defines its position during one of the four kinematic states. During animation, cranial elements simultaneously transition between these four states with the animation software smoothly interpolating between each of the four transformation matrices (Supplementary Video 2).

Because the μCT scanned individual was fixed in a state of suspensorial abduction and hyoid depression close to the maximum values observed in high-speed videos, the *in situ* fixed positions of the cranial elements were used for this later stage. Consequently, only three other kinematic stages needed to be recreated: resting-state, jaw opening prior to hyoid depression, and hyoid depression prior to jaw closing.

To recreate the resting-state position, each palate was fully adducted along its sliding articulation with the basipterygoid process, keeping its anterior process in articulation with the vomer. The hyomandibula, along with the dermal cheek, was rotated to rearticulate its connection with the palatoquadrate while maintaining articulation with the neurocranium. The jaws were rotated into fully closed position and the mandibulohyoid ligaments protracted to articulate with retroarticular process. The ceratohyals were protracted to maintain connection with the mandibulohyoid ligaments and repositioned to fit snuggly under the palate without intersecting the preopercles. The interhyals were then rotated to match the new orientation between the ceratohyals and the hyosymplectics.

To position the cranial elements into jaw opening prior to hyoid depression, the jaws were rotated from resting position into gape similar to the gape angles recorded in the high-speed videos. Some long-axis rotation of the jaws was necessary during jaw opening to prevent the medial flanges of the prearticular from intersecting with the ventrolateral wing of the palatoquadrate that lies anteromedial to the jaw joint. The mandibulohyoid ligament was retracted to match the new orientation of the retroarticular processes, and the ceratohyal was retracted to follow the mandibulohyoid ligament. The interhyal was then pivoted posteriorly in its articular socket with the hyosymplectic to follow the new position of the ceratohyal.

To recreate hyoid depression prior to jaw closing, the neurocranium was elevated to the peak angle seen in high-speed videos (10°, Table 1). Then the pectoral girdle was retracted by half the amount observed. By maintaining distance between the hypohyals and pectoral girdle, the hyoid was similarly rotated by approximately half its full observed value from high-speed videos. Prior to jaw closing, the jaws were set to their maximum value, which was only a slightly higher gape angle compared to the previous stage. Although the ceratohyals were depressed in this stage, the interhyals were kept at their maximum posterior rotation relative to the preopercle.

## 3 RESULTS

The feeding apparatus of *Atractosteus spatula* is capable of rapidly unfolding from a tightly compacted resting state to a massively expanded volume in less than 42 ms. High-speed videos document the timing of this expansion, which occurs in an anterior-to-posterior sequence and appears to generate a unidirectional flow of water through the feeding apparatus. The feeding apparatus of *A. spatula* functions as a cranial linkage system which binds many separate mobile elements into a single expansive mechanism using muscles and ligaments. Specialized joints, found throughout the feeding apparatus of *A. spatula* (visualized using μCT), increase the range of motion of the linkage system and enable the multiple configurations of the feeding apparatus seen during feeding. Control of this feeding system is modulated by a plesiomorphic complement of osteichthyan feeding musculature. Although unspecialized, many of these muscles nevertheless are found to serve dual roles within the feeding mechanism, necessary for driving the anterior-to-posterior expansion of the linkage mechanism seen in feeding videos.

### Kinematic timing during the feeding strike of Aractosteus spatula

#### Phase I: Jaw opening and lateral snapping

Jaw opening in *Atractosteus spatula* was rapid, lasting an average of 17.4 ± 0.7 ms (Table 1). As lateral axial bending swept the jaws into position around the prey item, the neurocranium elevated slightly, 10.2 ± 2.2° relative to the pectoral girdle and body (Table 1). As the lower jaws rotated ventrally, the ligamentous maxilla swung anteriorly and the medial wings of the prearticular flared laterally, exposing the unpigmented tissue underneath (Fig. 2e-h). Slight movements of the pectoral girdle were sometimes visible at this stage, although onset of pronounced pectoral girdle retraction did not begin until the next phase of the feeding strike.

During the initial stages of jaw opening, the floor of the buccal cavity could be seen buckling inwards, indicating negative pressure in the space between the expanding jaws prior to the initiation of hyoid depression (Fig. 2c). Although the prey item was held by the forceps, suction acted upon the floating portion of the prey item, drawing it towards the buccal cavity (Fig. 2). Lateral movements towards the prey continued until maximal gape (Fig. 3c), coinciding with the jaws positioned around the prey item. At this point, lateral movement of the jaws abruptly decelerated and the predator transitioned to anterior acceleration accompanied by hyoid depression (Figs. 2d-e and 3c).

#### Phase II: Hyoid depression and forward lunging

Hyoid depression was variable in timing and its onset ranged temporally between the onset of jaw opening and peak gape. This variation showed individual effect, with the two largest individuals in the study (A and B) regularly opening their jaws fully prior to hyoid depression (Fig. 2c), while the smallest individual (E) typically initiated hyoid depression and jaw opening at the same time (Fig. 3). On average, onset of hyoid depression was delayed an average of 9.1 ± 2.8 ms between individuals after initiation of jaw opening, at which point the jaws had already reached approximately half of peak gape (Fig. 3, Table 1).

The initiation of hyoid depression marked a distinct shift in the movement of cranial elements in high-speed videos. The folds of skin and musculature that were held in a constricted and elevated state in the initial phases of jaw opening began to distend ventrally to accommodate depression of the hyoid bars and basihyal toothplate (Fig. 2d). Pronounced rotation of the pectoral girdle resulted in a posteroventral displacement of the tip of the cleithrum, and the isthmus connecting the cleithrum and hyoid could be seen retracting from underneath the branchiostegal membrane (Fig. 2e-f). Lateral movement of the jaws decelerated, and anterior displacement toward the prey accelerated (Fig. 3).

Suction continued to draw the prey item into the oral cavity (Fig. 2d). Movement of the prey into the oral cavity occurred throughout hyoid depression, even while anterior acceleration and/or jaw closing might have been expected to displace the water containing the prey item out of the jaws.

#### Phase III: Jaw closing and suspensorial abduction

While jaw closing typically marks the onset of the “compressive phase” in other actinopterygians (Lauder, 1980a; Liem, 1978), in *Atractosteus spatula* jaw closing coincided with the greatest volumetric expansion of the oral cavity. Due to the late onset of hyoid depression, the vast majority of hyoid depression occurred after peak gape (Figs. 2 and 3). Furthermore, suspensorial abduction occurred exclusively during jaw closing (Table 1, Fig. 3). The jaws closed even as other elements – such as the cleithrum, suspensorium, and hyoid – continued to rotate and abduct, making jaw closing an “expansive” kinematic event. At the moment of jaw closure (an average of 23.3 ± 3.3 ms after peak gape), the feeding apparatus was fully expanded, coinciding with both peak hyoid depression and peak suspensorial abduction (Table 1, Fig. 3).

Suspensorial abduction resulted in a large posterior expansion of the feeding apparatus during this phase, laterally flaring the jaw joints, cheeks, and opercular flaps. Suspensorial abduction was evident posteriorly as the dermal cheeks rotated dorsolaterally along their hinge with the braincase, which resulted in 44.9 ± 7.6% lateral expansion of the cranium occurring at the opercular hinge by peak abduction (Table 1, Fig. 3). Anteriorly, the palate could be seen sliding laterally, exposing the underlying unpigmented skin of the ectopterygoid (Fig. 2e). This portion of the ectopterygoid is normally covered by a flanged portion of the frontal and premaxilla in the form of a scarf joint (see below) but was clearly visible during suspensorial abduction as a longitudinal white line running the length of the rostrum in dorsal view (Fig. 2e-h).

#### Phase IV: Compression and opercular abduction

Although the opercular flaps were laterally abducted during suspensorial abduction, distinct opercular abduction beyond movements of the cheeks was not apparent until the compressive phase of the feeding strike (Fig. 3), after the jaws had fully closed (Fig. 2f-h). During the reset phase, the hyoid elevated, the pectoral girdle protracted, the ventrolateral margins of the cheeks adducted, and the palate slid back beneath the frontal and premaxilla. This caused the opercular flaps to adduct medially as well; however, the dorsal portion of the opercular flaps were continually held open until all other elements were brought back into resting position, and water could be seen flowing posteriorly through the open gill covers. The prey item was held in the feeding apparatus for an extended period of time prior to deglutition as the hyoid elevated and suspensorium slowly adducted (Table 1, Fig. 3).

### Osteological components of the cranial linkage system

The feeding apparatus of *Atractosteus spatula* is a linkage mechanism composed of elements of the mandibular arch (palatoquadrate, dermal palate, and lower jaws), the hyoid arch (hyosymplectic, interhyal, and hyoid bar), and the pectoral girdle (supracleithrum and cleithrum), all suspended from the neurocranium and bound together into a single linkage system by muscles, ligaments, fibrous connective tissue, and synovial joints. Internal to each of the arches and pectoral girdle, specialized joints between individual elements increase the range of motion of each functional unit. The bones comprising these functional units have been described in detail for *Atractosteus* and *Lepisosteus spp.* in other studies (Arratia & Schultze, 1991; Grande, 2010; Jollie, 1984; Wiley, 1976), so the following descriptions focus on new information on soft-tissue anatomy documented through contrasted-enhanced μCT and *in situ* manipulations.

#### Mandibular arch

The palatoquadrate and dermal palate form three distinct articulations with the braincase. Anteriorly, a small forward-facing process of the dermopalatine fits into a socket formed dorsally by the premaxilla and ventrally by the vomer (Fig. 5e). This joint is filled with dense connective tissue, allowing only slight mediolateral movement at its articulation (Fig. 5e); however, it also corresponds with the anterior terminus of an elongate, longitudinal scarf joint that runs the length of the rostrum anterior to the orbits (Fig. 5f). This “rostral” scarf joint is formed by lateral flanges of the premaxilla and frontal bones overlapping the ectopterygoid, and it can be seen sliding laterally during feeding (Fig. 2e-h). Within this scarf joint, a pair of cartilaginous processes extend dorsolaterally from the ethmoid cartilage to support lateral flanges of the overlying frontal bones. These processes correspond to a thickened, anteriorly tapering portion of the palatoquadrate cartilage lying in a dorsal groove of the ectopterygoid, the pars autopalatina (*sensu* Konstantinidis et al., 2015) (Fig. 5c). This unusual ethmoid-autopalatine articulation allows mediolateral sliding, while restricting dorsolateral rotation of the palate. Posteriorly, the metapterygoid ossification of the palatoquadrate articulates with an enlarged basipterygoid process. This joint is unique in gars (Arratia & Schultze, 1991; Grande, 2010; Jollie, 1984; Wiley, 1976) and provides a broad expanded surface for mediolateral excursions of the metapterygoid. Unlike the hyosymplectic, movements of the palate pivot around a vertically oriented axis of rotation through the dermopalatal-vomerine (dpt-vo) joint (Table 2, Fig. 5e) (see below).

**Figure 5.**
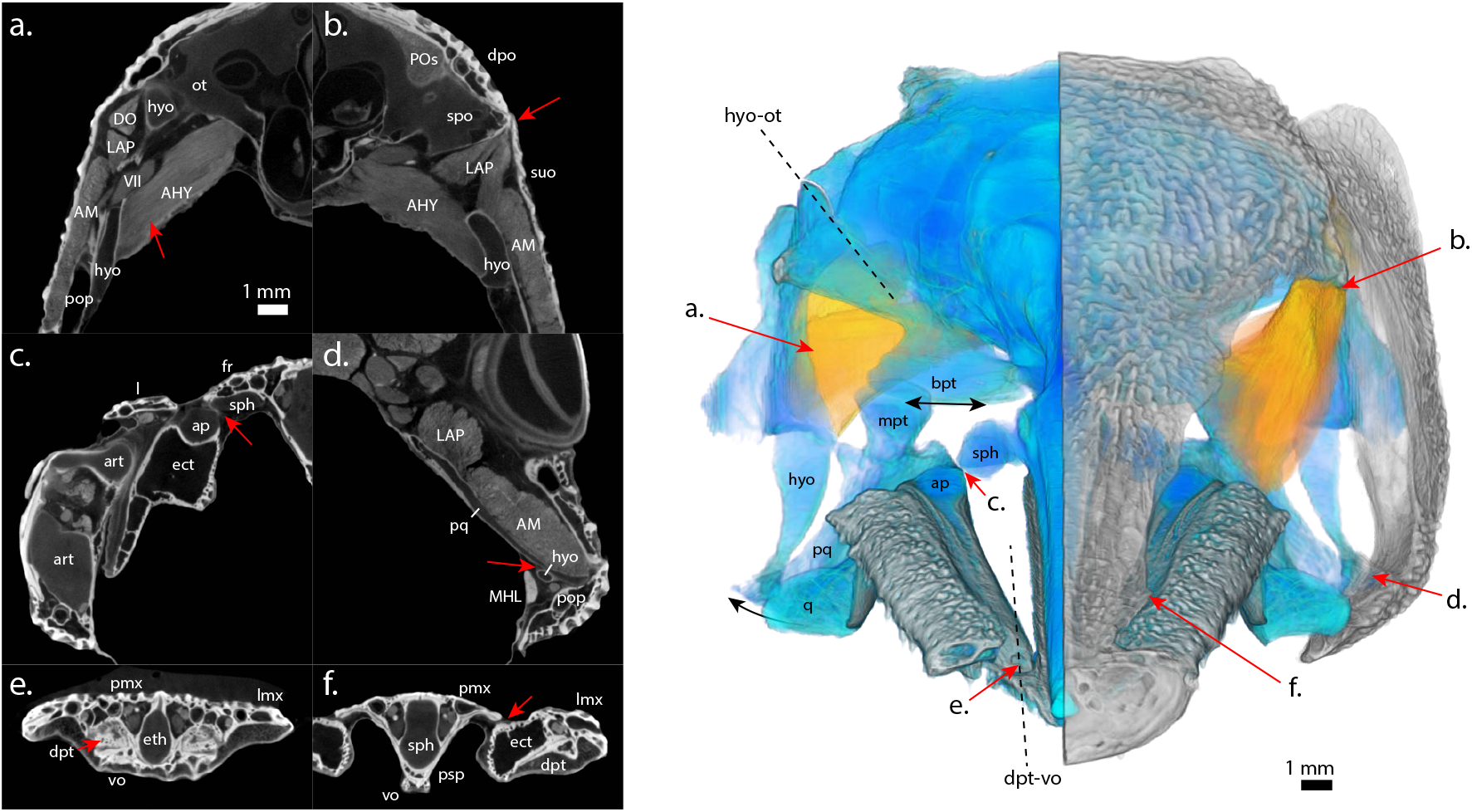
Components of the mechanism of cranial kinesis of *Atractosteus spatula*. Reconstructed cranial anatomy based on contrasted-enhanced μCT data, detailing the joints, bones, cartilages, muscles, and axes of rotation associated with a mobile suspensorium. (*a*) The hyosymplectic (hyo) has an oblique articular axis (hyo-ot) with the otic region of the braincase (ot). It is intimately connected to the medial surface of the preopercle (pop, the main component of the dermal cheek), and it has a broad attachment area of the adductor hyomandibulae (AHY) near the foramen for the hyomandibular trunk of facial nerve (VII). (*b*) The sphenotic (spo) is the origin for the levator arcus palatini (LAP), and it lies dorsal to a simple hinge line between the suborbitals (suo) and dermal braincase (e.g., dpo). (*c*) Smooth, cartilaginous processes of the pars autopalatina (ap) correspond with and enable sliding relative to specialized lateral projections of the sphenoid (sph). (*d*) A flexible intrasuspensorial joint is formed by a cartilaginous overlap of the posterior palatoquadrate (pq) and anterior hyosymplectic (hyo), enabling twisting between these two portions of suspensorium. (*e*) Rotation of the palate relative to the braincase occurs around a vertical axis running through the dermopalatine-vomerine joint (dpt-vo). (*f*) A broad, overlapping scarf joint between the premaxilla (pmx) and ectopterygoid (ect) lacks dense connective tissue and enables lateral sliding of the palate relative to the braincase (the white line that appears along the rostrum in dorsal view of high-speed videos, Fig. 2*e-h*). For abbreviations, see Table 2.

Specialized joints of the lower jaws of *Atractosteus spatula* correspond with mediolateral movements of the palate. As reported for *Lepisosteus* (Askary et al., 2016; Haines, 1942), the quadrate-articular joint of *A. spatula* is synovial, and the convex, rounded condyle of the quadrate allows a wide range of dorsoventral as well as mediolateral motion. This joint is exclusively suspended from the palatoquadrate without any support from the hyoid arch (Grande, 2010; Wiley, 1976) (see below). In contrast to the condition in *Lepisosteus* (Grande, 2010), the mandibular symphysis of *A. spatula* is short and flexible, allowing mediolateral wish-boning of the lower jaws. In gars, the symphysis does not contain Meckel’s cartilage (Grande, 2010; Jollie, 1984). Instead, in *A. spatula*, it is filled with cartilage that appears to be continuous with and of a similar composition as the rostrohyal (*sensu* Hilton, Konstantinidis, Schnell, & Dillman, 2014), forming a thin, anterior buccal floor (Fig. 6).

**Figure 6.**
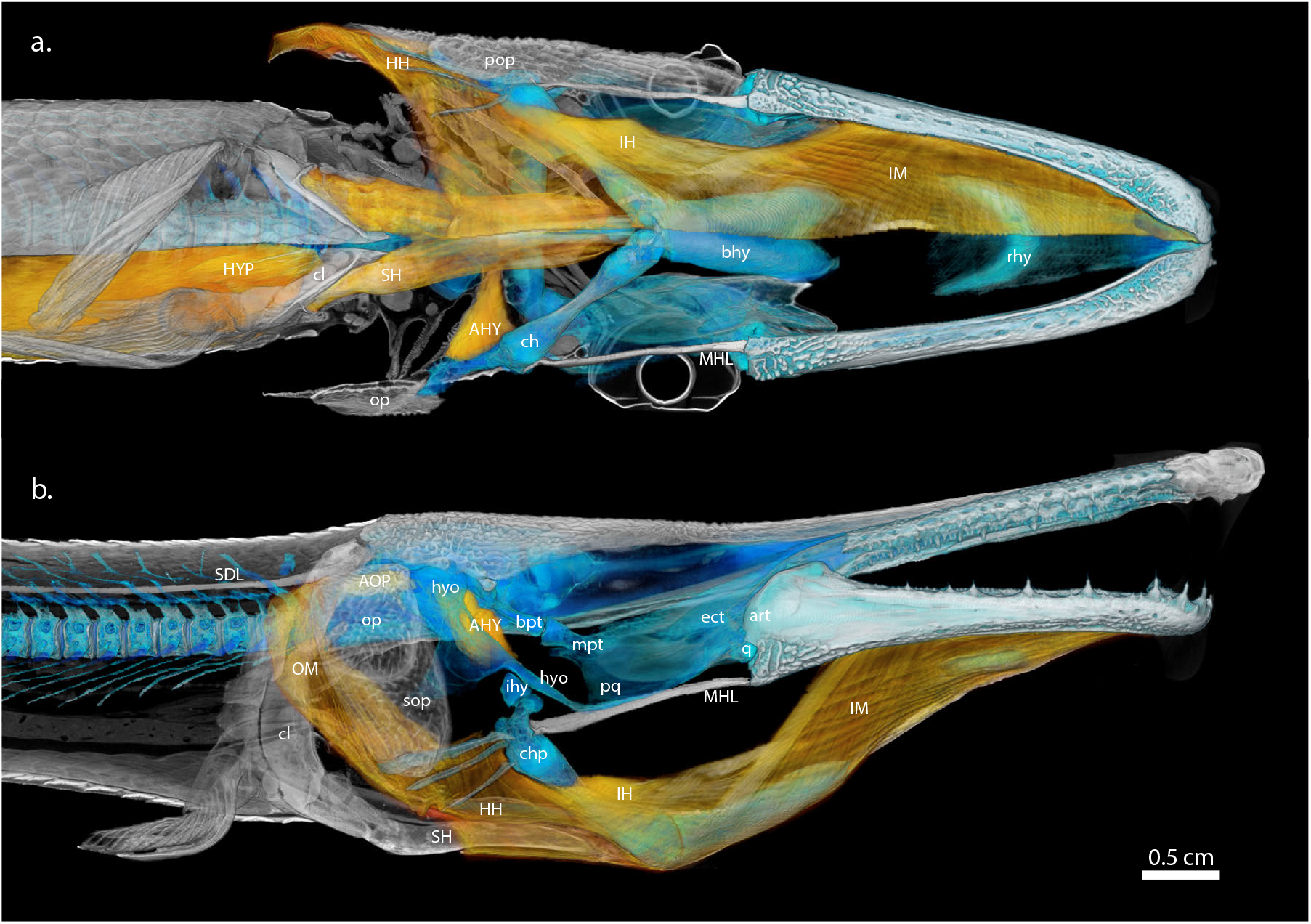
Fully expanded feeding apparatus of *Atractosteus spatula* detailing the hyoid constrictor sheath and underlying skeletal elements. Reconstructed cranial anatomy based on contrast-enhanced μCT data of a feeding mechanism fixed in expanded position (see Fig. 4*d+h*) in (*a*) ventral and (*b*) right lateral views. Shown are the superficial hyoid constrictor muscles, with the dermal cheek and underlying mandibular constrictor musculature removed. Ventral hyoid constrictors (IM, IH, HH) wrap around the sternohyoideus (SH) and hyoid apparatus (chp, bhy), while dorsal hyoid constrictors (AHY, AOP) span between the operculum (op), hyosymplectic (hyo), and the braincase. At rest, these muscles are in position to restrict rotation of the hyoid apparatus and abduction of the cheeks during the earliest phases of the feeding strike but still permit hyoid retraction via the SH, opening the jaws by means of the mandibulohyoid ligament (MHL). During later phases of the feeding strike, these muscles relax to allow volumetric expansion of the feeding apparatus (shown here). See Fig. 7 for dorsal view of the adductor operculi (AOP). For abbreviations, see Table 2.

Gars have a distinct retroarticular process ventral to the jaw joint where the mandibulohyoid ligament attaches (Fig. 6). There is no evidence for an interoperculomandibular ligament in *Atractosteus spatula*, despite the clear presence of other ligaments in contrast-enhanced μCT (Fig. 5d). No other ligament or muscle attaches to the retroarticular process other than the mandibulohyoid ligament, which makes hyoid retraction due to sternohyoideus contraction the sole input responsible for depression of the lower jaws (Lauder, 1980a). Due to the close insertion of the mandibulohyoid ligament to the jaw joint (Kammerer et al., 2006), only slight posterior movements of the hyoid are necessary to open the jaws fully.

#### Hyoid arch

The hyosymplectic is suspended from the spheno-pterotic ridge (*sensu* Allis, 1919) of the auditory capsule, directly ventral to the dermopterotic (Grande, 2010). In the ontogeny of gars the hyosymplectic fuses to the medial surface of the preopercle (Grande, 2010), and its movements are directly linked to movements of the dermal cheek. Contact between the hyosymplectic and palatoquadrate is limited to a small overlap between the posteroventral angle of the palatoquadrate cartilage and the anterior portion of the symplectic (Grande, 2010) (Figs. 5d and 6). Gars are unusual in that the hyoid arch does not contact any ossifications of the palatoquadrate directly (Arratia & Schultze, 1991; Grande, 2010; Wiley, 1976). Unlike in *Amia*, there is no direct contact between the metapterygoid and the hyosymplectic (Arratia & Schultze, 1991; Grande, 2010; Wiley, 1976) (Fig. 6). As a result, it is possible for hyosymplectic-palatoquadrate joint to twist during suspensorial abduction. As the palate slides mediolaterally along the metapterygoid-basipterygoid joint, the hyosymplectic-cheek complex rotates dorsolaterally (Fig. 5, Supplementary Video 2). Rotation of the hyosymplectic-cheek complex is centered around a horizontal, anteromedially oriented axis of rotation through the hyomandibular-otic (hyo-ot) joint (Table 2, Fig. 5). Unlike the “rostral” scarf joint (see above), the joint between the dermal cheek and skull roof is a simple hinge which remains articulated during suspensorial abduction (Fig. 2e-h).

The interhyal rests in a synovial joint on the medial wall of the preopercle, bounded dorsally by the hyosymplectic (Fig. S1). In contrast-enhanced μCT, this joint appears similar in cross-section to the quadrate-articular joint, which is a known synovial joint (Askary et al., 2016; Haines, 1942) (Fig. S2). The interhyal is capable of a wide range of motion upon the dorsal process of the posterior ceratohyal, which is large, rounded, and acts as a ball-in-socket joint.

Due to the mobility of the interhyal-ceratohyal joint and the hypohyal-basihyal joint, the hyoid bar is capable of retraction relative to the hyosymplectic, rotation relative to the interhyal, and abduction relative to the sagittal plane. The mandibulohyoid ligament originates on the neck of the dorsal process of the posterior ceratohyal (Fig. S1), close to the point of rotation of the ceratohyal-interhyal joint and attaches to the retroarticular process of the lower jaws. The mandibulohyoid ligament links hyoid retraction with jaw depression (Lauder, 1980a), but it also causes hyoid protraction when the jaws are adducted. The sternohyoideus attaches to the hypohyals of the hyoid bar and serves to link movements of the hyoid apparatus and pectoral girdle.

#### Pectoral girdle

The supracleithrum attaches to the back of the dermatocranium by means of a small cup-shaped articulation that extends anterodorsally to the lateral extrascapular and posttemporal (Grande, 2010). The supracleithrum is bound to the dermatocranium by connective tissue through which the lateral line passes (Grande, 2010). As a result, movement at this joint appears restricted, although the neurocranium is capable of slight elevation relative to pectoral girdle.

In contrast, the cleithrum is only loosely connected to the supracleithrum by connective tissue and a small slip of the cucullaris muscle originating on the medial surface of the supracleithrum (Fig. S3). The cleithrum is capable of large posteroventral rotation relative to the supracleithrum, and is easily disarticulated during manual manipulation. The cleithrum is the origin of the sternohyoideus, which acts as a muscular link between the pectoral girdle and hyoid arch. The cleithrum is also the attachment point for powerful hypaxial musculature which drives the expansion of the cranial system due to posteroventral retraction of the sternohyoideus (see results: hypaxials).

#### Cranial Linkage

The interconnections of all these elements can be demonstrated on a fresh *Atractosteus spatula* specimen via *in situ* manipulations (Fig. 4, Supplementary Video 1). By simply retracting the pectoral girdle, simulating input of the sternohyoideus and hypaxial musculature, the entire feeding system expands: the hyoid depresses, the cheek-hyosymplectic rotates dorsolaterally, the palate slides laterally, and the jaws weakly open (Fig. 4c). Counterintuitively, closing the jaws in this state causes the cheek and suspensorium to expand further laterally, whereas pressing the cheeks together cause the jaws to open wider (Supplementary Video 1). Maximal jaw opening occurs during manipulation with the cheeks adducted and the hyoid held in an elevated position (Fig. 4b, Supplementary Video 1). This demonstrates that additional muscular input is needed – other than sternohyoideus and axial musculature – to produce the multiple configurations of the feeding apparatus seen in the high-speed videos throughout the feeding strike.

### Muscular input to the feeding mechanism

Manipulation of fresh specimens indicates that additional muscular input other than the sternohyoideus is necessary to create the multiple configurations of the feeding apparatus seen in high-speed videos (Fig. 4, Supplementary Video 1). Gars maintain only a plesiomorphic complement of osteichthyan feeding musculature with which to modulate the anterior-to-posterior expansion of their cranial linkage system. However, many of these muscles appear to have secondary functions typically found in more specialized muscles of derived actinopterygians. Emphasis is placed here on muscles with atypical morphology and potentially multiple roles as a result of cranial expansion during the feeding strike of *Atractosteus spatula*.

#### Sternohyoideus

The sternohyoideus connects the cleithrum to the hypohyals and basihyal toothplate (Figs. 6 and 7). This muscle is responsible for both hyoid retraction and rotation, which in gar are two separate kinematic events (Figs. 2 and 3). More than any other muscle, its function changes during the feeding strike due to repositioning of its origin and insertion. At rest, this muscle passes just below the point of rotation of the interhyal-ceratohyal joint (Fig. 8). During early activity, the sternohyoideus serves primarily to retract the hyoid. The ventral isthmus containing the sternohyoideus passes dorsally to and is separate from the ventral hyoid constrictor musculature (Farina et al. 2015), which allows the sternohyoideus to retract freely relative to those muscles. However, as the pectoral girdle rotates posteroventrally and the ventral hyoid constrictors relax, this line action falls well below the point of rotation of the interhyal-ceratohyal joint (Fig. 7 and 8). As a result, activity of the sternohyoideus in the later stages of the feeding strike results in rotation rather than retraction of the hyoid, and the interhyal and dorsal process of the posterior ceratohyal are able to protract as the hyoid bar rotates ventrally during jaw closing (Fig. 8d).

**Figure 7.**
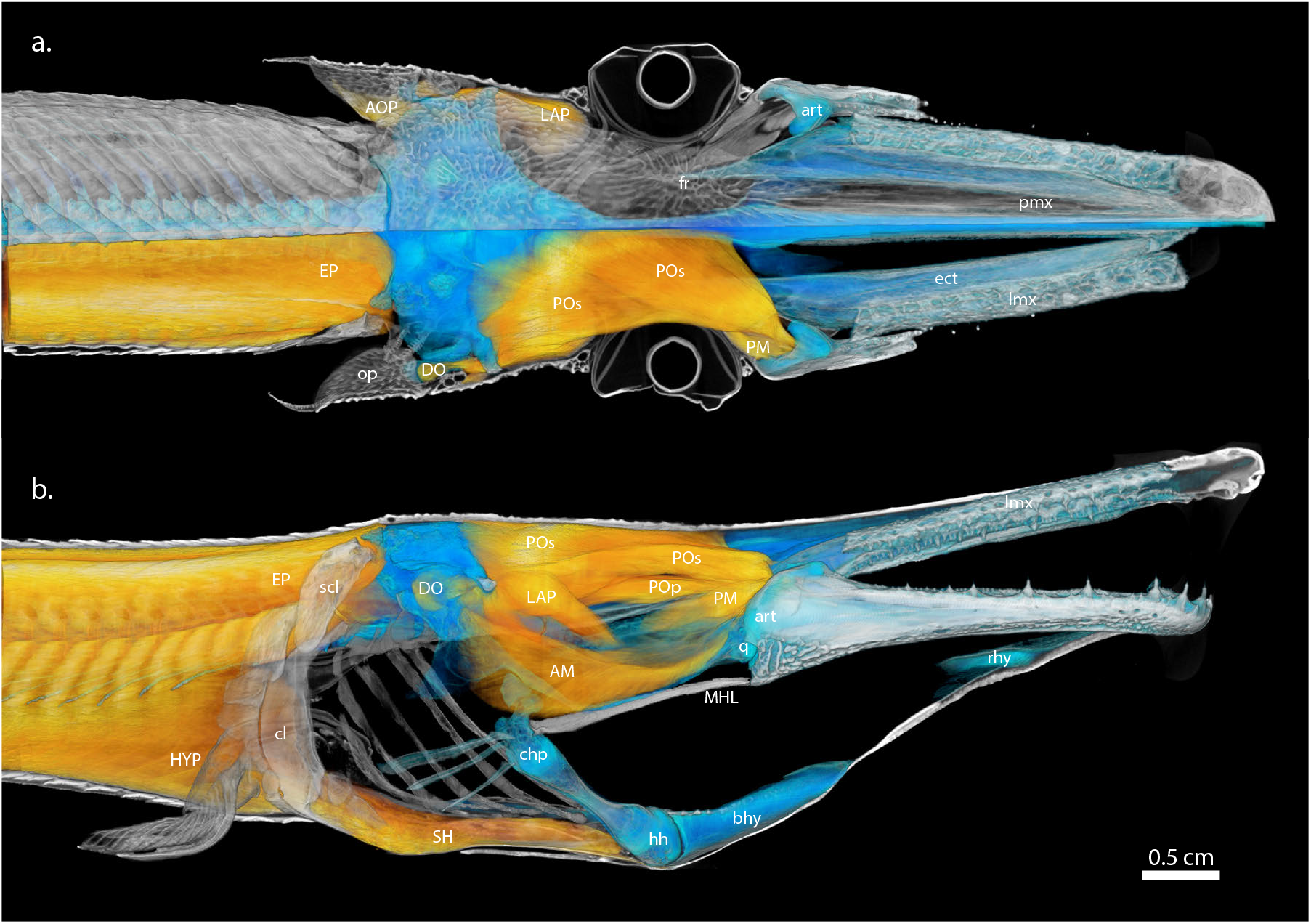
Fully expanded feeding apparatus of *Atractosteus spatula* detailing axial musculature and constrictors of the mandibular arch. Reconstructed cranial anatomy based on contrast-enhanced μCT data of a feeding mechanism fixed in expanded position (see Fig. 4*d+g*) in (*a*) dorsal and (*b*) right lateral views. Superficial elements, such as the dermal cheek, operculum, scales, and hyoid constrictor musculature have been removed (see Fig. 6). Epaxials (EP) and hypaxials (HYP) are responsible for elevating the neurocranium and retracting the pectoral girdle (cl), which vertically raises the interhyal-ceratohyal (chp) joint above the line-of-action of the sternohyoideus (SH). This shifts the function of the sternohyoideus (SH) from hyoid retraction (jaw opening) to hyoid rotation (pharyngeal expansion). Lateral abduction of the cheek and palate is mediated by the levator arcus palatini (LAP) but also assisted by the jaw adductors. Jaw adduction protracts the interhyal-ceratohyal joint via the mandibulohyoid ligament (MHL), holding the posterior ceratohyals in place anteriorly, while the sternohyoideus (SH) posteroventrally rotates the hypohyals (hh), causing the hyoid bars to depress ventrally and rotate laterally, flaring of the cheeks. For abbreviations, see Table 2.

**Figure 8.**
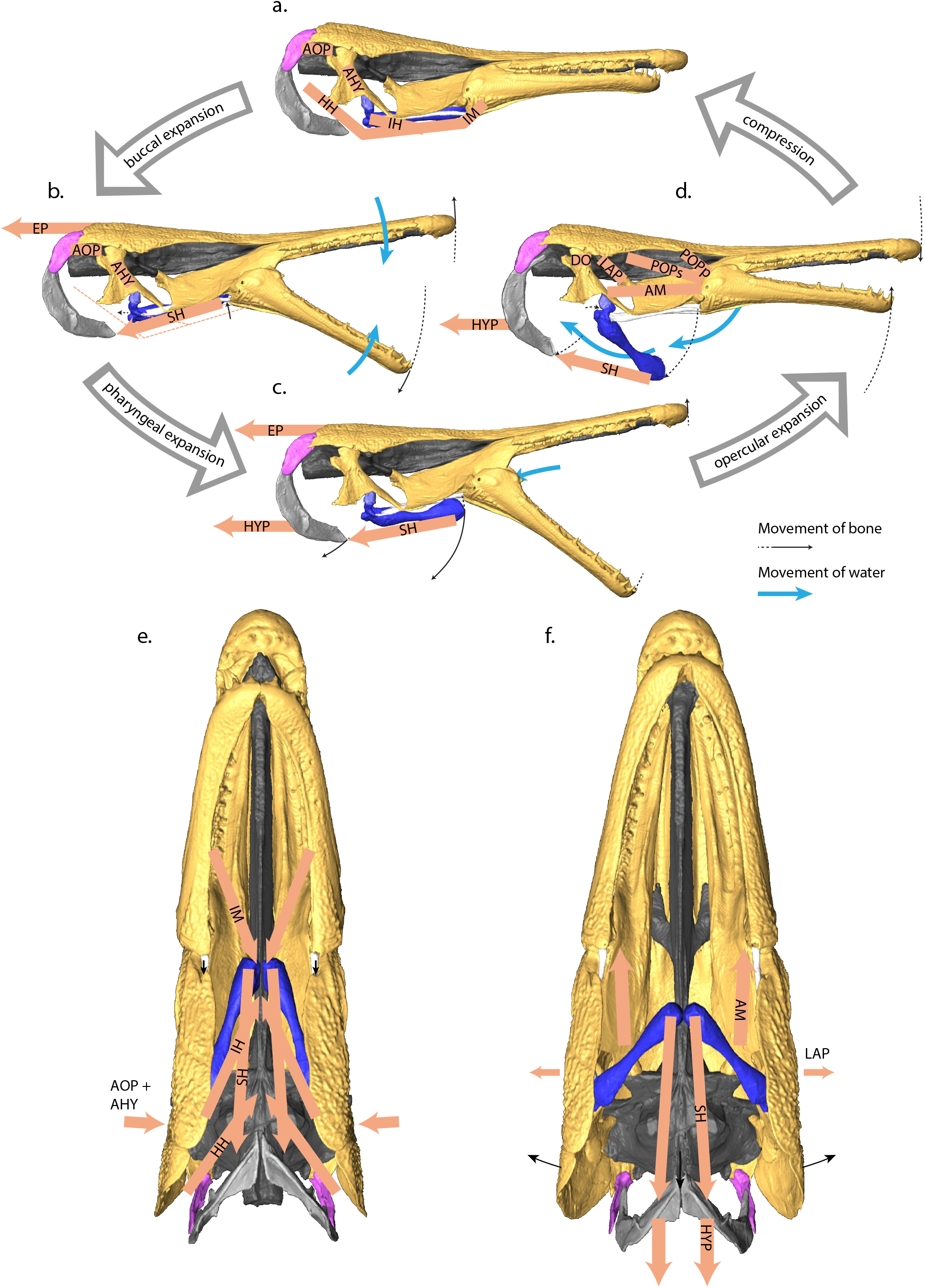
Reconstructed gape cycle and feeding kinematics of *Atractosteus spatula*. (*a-d*) The feeding apparatus of alligator gars is capable of successive expansions of the buccal, pharyngeal, and opercular cavities, modulated by hyoid constrictor musculature, which causes an anterior-to-posterior flow of water through the feeding apparatus (blue arrows). (*a*) At resting state, the ceratohyals and sternohyoideus are positioned within a sheath of hyoid constrictor musculature (IM, IH, HH, AOP, AHY). (*b*) Buccal expansion is caused by posterior retraction of the ceratohyals by the sternohyoideus (SH) while hyoid elevation is maintained by ventral hyoid constrictors (IM, IH, and HH) and ceratohyal adduction is maintained by dorsal hyoid constrictor musculature (AOP, AHY). (*c*) Pharyngeal expansion is enabled by relaxation of the ventral hyoid constrictors, which allows the sternohyoideus to ventrally rotate the ceratohyals as the pectoral girdle begins to rotate and depress the line-of-action of the sternohyoideus. (*d*) Opercular expansion is caused by further rotation of the pectoral girdle by hypaxial musculature while the jaws close due to jaw adductors (AM). (*e*) Suspensorial adduction is maintained in the early stages of jaw opening due to the hyoid constrictors. (*f*) Opercular expansion is the combined result of lateral rotation of the suspensorium and hyoid bars during jaw closing, facilitated by jaw adductors (AM, POPs, POPp) and levator arcus palatini (LAP). For abbreviations, see Table 2.

To serve as both rotator and retractor of the hyoid, the sternohyoideus has an atypical morphology. In *Polypterus* and *Amia*, the sternohyoideus is divided into three muscle blocks by two transverse septa (Allis Jr, 1922; Edgeworth, 1935; Lauder, 1980a), but in *Atractosteus spatula*, these muscle blocks are deeply nested within each other (Fig. S4). This pennate morphology suggests the muscle is optimized for force transmission rather than displacement. To open the jaws fully requires very little displacement of the retroarticular process due to the low mechanical advantage of the jaws (Kammerer et al., 2006); however, hyoid rotation requires far more displacement than the sternohyoideus itself appears capable of (Fig. 2). Due to its structure, the sternohyoideus is able to convey powerful posterior retracting forces to the hyoid over a short distance, but any further retraction of the hyoid is likely conveyed through the sternohyoideus by hypaxial-driven pectoral girdle rotation (see below).

#### Hyoid constrictors

An extensive, continuous sheath of hyoid constrictor musculature supports the ventral floor of the buccal, pharyngeal, and opercular cavities. From anterior to posterior, this muscular sheet is composed of the intermandibularis (originating from the paired Meckel’s cartilages), interhyoideus (originating from the anterior ceratohyals), and hyohyoideus (inferioris/superioris) (spanning between the branchiostegal rays and originating from the medial surfaces of the subopercle). These muscles intermingle fibers and join together at the midline to form a muscular sling that ventrally supports the rostrohyal, basihyal, hyoid, and sternohyoideus (Fig. 6). The ventral constrictor sheet functions primarily to return the hyoid apparatus to its resting state after the feeding strike is complete (interhyoideus = protractor hyoideus *sensu* Wiley, 1976), and these muscles are capable of extreme distension during hyoid depression to accommodate the rotated hyoid (Figs. 2 and 6). However, these muscles also appear to maintain hyoid elevation during the earliest phases of the feeding strike (Figs. 2c and 4b). By constraining action of the sternohyoideus to only allow posterior retraction of the hyoid, these muscles may have an important secondary role in jaw opening (see discussion).

A band of dorsal hyoid constrictors binds the dorsal and posterior edge of the opercular flap. These muscles include the adductor hyomandibulae (originating from the lateral wall of the otic capsule and inserting onto the medial surface of the hyosymplectic) and the adductor operculi (originating from the connective fascia just lateral to the otic capsule and inserting onto the medial surface of the opercle). Unlike *Lepisosteus* (Konstantinidis et al., 2015; Lauder, 1980a), no distinct adductor arcus palatini muscle could be identified in *Atractosteus spatula*: the adductor hyomandibulae does not have any fibers that insert onto the palatoquadrate (Fig. 6). Like *Lepisosteus* (Konstantinidis et al., 2015; Lauder, 1980a), the adductor operculi shows no sign of subdivision, and no specialized jaw opening muscle corresponding to the levator operculi in *Amia* could be identified. Instead, the adductor operculi intermingles fibers with the opercular membrane, which also subsequently fuses with the hyohyoideus superior to form a complete opercular valve around the margins of the opercular flap. The primary function of these muscles is to adduct of the operculum, dermal cheek, and suspensorium at the end of the feeding strike (Fig. 2f-h). However, these muscles also appear to maintain suspensorial and opercular adduction in the earliest phases of the feeding strike (Fig. 2a-d). As with the ventral hyoid constrictors, by restricting action of the sternohyoideus to hyoid retraction rather than rotation (in this case laterally), dorsal hyoid constrictors may have an important secondary role in jaw opening as well (see discussion).

#### Epaxials

Epaxial muscles attached to the back of the neurocranium are powerful levators of the neurocranium. These muscles surround the neural spines of the vertebral column and a well-developed supradorsal ligament (Fig. 6b). While neurocranial elevation is known to contribute to jaw opening in teleosts (Liem, 1978), with the lower jaws suspended entirely from the palatoquadrate (see above) (Grande, 2010; Wiley, 1976), it is less clear how this elevation could contribute to rapidly increasing gape angle in gars, and neurocranial elevation persists well into jaw closing when the hyoid is depressed (Fig. 3). However, by elevating the hyoid while the ventral hyoid constrictors are active, the epaxial musculature may nevertheless contribute to jaw opening (see discussion).

#### Hypaxials

While it is recognized that hypaxial musculature functions to stabilize the pectoral girdle during jaw opening in gars (Lauder, 1980a), hypaxial musculature attaching to the posterior portion of the cleithrum also drives the posterolateral rotation of the shoulder girdle (Fig. 4d+h). The cleithrum rotates extensively during the later stages of the feeding strike of *Atractosteus spatula* (Figs. 2 and 3), and this rotation marks a kinematic shift in the feeding strike of *A. spatula* that coincides with the transition between hyoid retraction and hyoid rotation as well as the transition from flat-plate suction to hyoid-based suction (Fig. 2b-d). By altering the line-of-action of the sternohyoideus and enabling the posteroventral displacement necessary for hyoid depression (see above) (Fig. 4d), hypaxial musculature appears to play an important role in expanding the feeding system in the later stages of the feeding strike (Figs. 2d-f, 4d+h).

#### Adductor mandibulae

The divisions of adductor mandibulae function primarily as jaw adductors during jaw closing. These muscles include the preorbitalis muscles, which originate on the neurocranium, as well as the adductor mandibulae and palatomandibularis muscles, which originate from the palate, preopercle and hyosymplectic. Attachments of jaw adductors to the mandible can be categorized into at least three attachment areas (Fig. S2). The first is a broad attachment zone dorsolateral to the adductor chamber that provides insertion points for the adductor mandibulae tendon, the dorsal subdivision of preorbitalis superficialis, and the superficial palatomandibularis minor muscle (inserting just inside the adductor chamber) (Fig. S2a). The next attachment zone lies deep within the adductor chamber, distal to the jaw joint, providing an insertion for preorbitalis superficialis (and perhaps some intermingled fibers of profundis) (Fig. S2b). The last attachment zone lies deep within the adductor chamber, proximal to the jaw joint, providing insertions for palatomandibularis major and a tendinous slip of the preorbitalis profundus (Fig. S2c). While primarily responsible for closing the jaws, due to the mandibulohyoid ligament attachment to the posterior ceratohyal, jaw adductors also appear to have an indirect and secondary function in protracting the dorsolateral portion of the hyoid bar during jaw closing (Fig. 4h, Supplementary Video 1).

#### Levator arcus palatini

Levator arcus palatini (LAP) originates from the lateral most portion of the neurocranium, the sphenotic, and envelops the basipterygoid-metapterygoid joint where it inserts onto the dorsal and ventral surfaces of the metapterygoid. In this configuration it is able to act as an abductor of the palate relative to the braincase. However, LAP also originates partially on the suborbital bones of the cheek, although to a lesser degree than the condition reported for *Lepisosteus* (Lauder, 1980a). LAP also has small subdivisions extending posteriorly, the protractor hyomandibulae *sensu* Konstantinidis (2015), and a portion extending directly between the metapterygoid and the hyosymplectic (Fig. S5), which in *Lepisosteus* is reported to be entirely ligamentous (Arratia & Schultze, 1991). With these muscular connections between the palatoquadrate and hyosymplectic-cheek, it is possible that LAP serves to conjoin movements of the palatoquadrate and hyosymplectic, which are particularly flexible relative to each other (see above).

#### Dilatator operculi

The dilatator operculi (Fig. 7) is small and dwarfed by its functional antagonist, the adductor operculi muscle (Fig. 6). Opercular pumping during respiration might largely be due to suspensorial abduction/adduction rather than alternation of dilatator and adductor operculi activity. This suggestion is supported by observations of heavy respiration of gar in the lab, where suspensorial movements drive mediolateral movements of the operculum (personal observation).

## 4 DISCUSSION

The extent to which suction acted on the prey item during the feeding strike of *Atractosteus spatula* was an unexpected result of this study, but here we discuss how the need to maintain an effective suction generating mechanism may have influenced and constrained the evolution of feeding kinematics, cranial morphology, and muscular function in gars. While gars are known for their characteristic, lateral-snapping method of prey capture, *A. spatula* engages in a synergistic combination of jaw-ram and suction-feeding behaviors during the feeding strike that facilitates this feeding strategy. Suction generation relies on expansive cranial elements, and the skull of *A. spatula* has numerous modifications that maintain cranial kinesis despite modifications for lateral-snapping. Finally, plesiomorphic musculature modulates the timing of an anterior-to-posterior expansion of the feeding system of alligator gars, helping to explain the unusual firing patterns of lepisosteid cranial muscles during feeding, as well as demonstrating how hyoid constrictor musculature evolved specialized jaw opening roles in other taxa.

### Combined jaw-ram and suction in the feeding strike of Atractosteus spatula

It was previously thought that gars only used suction during prey processing and respiration (Porter & Motta, 2004; Werth, 2006), but this study shows that *Atractosteus spatula* utilizes suction during prey capture as well. Suction appears to act on the prey throughout all phases of the feeding strike of *A. spatula* to create a unidirectional flow of water into the oral aperture, complimenting the unique, lateral-snapping feeding strategy of gars. Feeding kinematics of *A. spatula* are largely similar to those reported for *Lepisosteus oculatus* and *L. platyrhinchus* in both timing and pattern (Lauder, 1980a; Porter & Motta, 2004), but high-speed video of *A. spatula* feeding indicates these kinematic events are also used to control the flow of water during the feeding strike. It is possible that many of the differences between gars and other, piscivorous, ram-feeding specialists (Porter & Motta, 2004) can be explained by a synergistic combination of lateral-snapping and suction generation in gars.

In the earliest stages of the feeding strike, alligator gars generate flat-plate suction, which appears to counteract the negative effects of a bow-wave during lateral sweeping. One of the problems of feeding in water, whether by ram or suction, is that prey approach can generate substantial bow-wave-induced disturbances (Holzman & Wainwright, 2009). Gars use surprise and rapid lateral acceleration to overcome prey (Porter & Motta, 2004), but that behavior would still incur a proportionately large bow-wave (Holzman & Wainwright, 2009), which has the potential to alert or even displace elusive prey (Day, Higham, Holzman, & Van Wassenbergh, 2015; van Leeuwen & Muller, 1984). High-speed videos of *Atractosteus spatula* feeding show no indication of a bow-wave pushing the prey item away, however, and instead it is drawn towards the space between the opening jaws even prior to hyoid depression (Fig. 2b). It is likely the jaws of *A. spatula* act as diverging flat-plates during jaw opening, which is known to generate suction in other biological systems (Vogel, 1994). While forward-lunging with broad-flat plates would be energetically costly due to the increased drag profile (Porter & Motta, 2004; Vogel, 1994), lateral-snapping with broad-flat plates appears to have some benefit for prey capture due to added suction generation.

Reduction of the drag is a possible explanation for the observed delay in hyoid depression seen in gars. Forward-lunging, body-ram predators often use pharyngeal and opercular expansion to avoid pushing prey away during prey approach (van Leeuwen & Muller, 1984), but those same expansions would increase drag for a lateral-snapping predator, such as gars. The platyrostral morphology of gars is believed to maintain a streamlined profile during lateral-snapping (Porter & Motta, 2004); however, in *Atractosteus spatula*, that streamlining exists only prior to hyoid depression. The buccopharyngeal floor of *A. spatula* is substantially distended during hyoid depression, and this coincides with rapid lateral deceleration (Fig. 3c). Pronounced delay of hyoid depression is noted in *Lepisosteus* species as well (Lauder & Norton, 1980; Porter & Motta, 2004), and it is likely that delayed hyoid depression is a strategy shared by all gar species for reducing drag during the lateral-snapping portion of the feeding strike. Additionally, in *A. spatula*, this delay appears to have the added beneficial effect of also saving hyoid-based suction for when it would be most effective during prey capture.

It has been argued that the primary function of ram behavior is to move the oral aperture close enough to the prey for suction to be effective (Wainwright et al., 2001), and the feeding kinematics of *Atractosteus spatula* surprisingly support this suggestion. Effective suction drops precipitously beyond a distance of one mouth diameter (Day, Higham, Cheer, & Wainwright, 2005; Day et al., 2015; Wainwright et al., 2001; Wainwright et al., 2015). In gars, the oral aperture is positioned at the base of the jaws, formed by the lateral wings of the ectopterygoids, medial wings of the prearticulars, parasphenoid, and basihyal toothplate. During lateral-snapping, the upper and lower jaws are swung around the prey item, but hyoid depression typically does not begin until the prey item is passing between the elongate tooth rows, anterior to the oral aperture. When this happens, feeding kinematics shift to forward lunging, pectoral girdle retraction, and hyoid depression, with the prey item being drawn further into the oral aperture even as the jaws begin to close (Fig. 2d).

Suspensorial and opercular abduction during jaw closure completes an anterior-to-posterior wave of expansion of cranial elements in *Atractosteus spatula* (Fig. 3a), and precisely timed anterior-to-posterior expansion is recognized as an important component of specialized suction feeding in actinopterygian fishes (Wainwright et al., 2015). In the case of *A. spatula*, suspensorial and opercular abduction occur exclusively during jaw closure and likely aide in directing fluid posteriorly as a counter to flat-plate compression, which might otherwise push the prey item out of reach of the jaws.

A central conclusion of this study is that lateral-snapping and suction operate synergistically to achieve prey capture in the feeding strikes of alligator gar. By simultaneously mitigating the bow-wave, reducing drag, and precisely timing the expansion of cranial elements for when they are most effective – all within 42 ms – *Atractosteus spatula* demonstrates one of the many ways jaw-ram and suction can be complementary feeding strategies. While gars are incapable of the jaw protrusion seen in many jaw-ram specialists (Porter & Motta, 2004), their strategy of laterally sweeping the jaws over prey can be considered a form of jaw-ram. This paper joins a growing body of literature recognizing the potential synergism and overlapping feeding strategies that jaw-ram and suction share (Cooper, Carter, Conith, Rice, & Westneat, 2017; Longo, McGee, Oufiero, Waltzek, & Wainwright, 2016).

### Specialized joints enable expansive capabilities of a platyrostral cranial linkage system

*Atractosteus spatula* maintains the expansive capabilities of its feeding apparatus despite dorsoventral compression of the cranium. While gars are noted for evolving elongate, platyrostral jaws associated with lateral-snapping (Porter & Motta, 2004), these same modifications place significant barriers on the ability to produce suction within traditional models of teleost suction feeding (Day et al., 2015; Lauder, 1980a; Liem, 1978; Muller, 1989; Muller, Osse, & Verhagen, 1982; van Leeuwen & Muller, 1984; Wainwright et al., 2015). In particular, restricted dorsolateral rotation of the palate (Lauder, 1980a), reduced hyoid to jaw length ratio (Muller, 1989), and relative reduction of the operculum (Hutchinson, 1973; Muller et al., 1982) were believed to be problematic for suction generation. There appears to be direct tradeoffs between effective suction generation and enhanced performance during ram-feeding (Norton & Brainerd, 1993); however, *A. spatula* demonstrates that suction generation can be maintained despite ram-related modifications to the feeding system through the use of specialized joints that maintain a compact, streamlined profile during lateral-snapping but allow rapid volumetric expansion for suction generation.

First, a specialized joint between the hyosymplectic and palatoquadrate portions of the suspensorium (Fig. 5d) increases the range of motion of the palate by freeing it to rotate independently of the long axis of the suspensorium. Intrasuspensorial mobility is required for rotation of any element of the suspensorium relative to other elements, and this decoupling enables the cheeks to rotate dorsolaterally while the palate slides mediolaterally. Flexible joints within the suspensorium have been identified in multiple groups of derived actinopterygians, which require separate movements of suspensorial elements. While typically these joints enhance the extreme jaw protrusion seen in jaw-ram specialists, such as the sling-jaw wrasse (Westneat & Wainwright, 1989), several long-jawed chaetodontid species (Ferry-Graham, Wainwright, Hulsey, & Bellwood, 2001), and several cichlid species (Waltzek & Wainwright, 2003), they have also evolved within the dorsoventrally compressed crania of benthic fish that utilize suction to capture prey, such as catfish (Arratia, 1990) and sturgeons (Carroll & Wainwright, 2003). It is likely that intrasuspensorial mobility is more commonplace than previously thought and could be used to enhance the range of suspensorial movements within their respective cranial linkages. Among gars, this enhanced range of motion of the cheek and palate enables gars to maintain suspensorial abduction despite dorsoventral compression of the cranium.

Additional specialized joints in the feeding apparatus of *Atractosteus spatula* directly facilitate its unique form of suspensorial abduction. The sliding “rostral” scarf joint between the palate and dermal skull roof (Fig. 5f), the gliding cartilages between the ethmoid and pars autopalatina (Fig. 5c), and the enlarged basipterygoid-metapterygoid articulation (Jollie, 1984) (Fig. 5), are derived features that support mediolateral rotation of the enlarged palate relative to the cheek. Maintained suspensorial abduction within the cranial linkage system of gars during the shift towards biting-based prey capture, likely provided the selective pressure for these specialized joints to evolve. Mobile joints between the braincase and palate persist despite significant restructuring of the palate and elongation of braincase in gars (Arratia & Schultze, 1991; Wiley, 1976).

In order to understand the function of suspensorial abduction we need to understand the limitations of the opercular pumping mechanism in dorsoventrally compressed taxa. The role of opercular abduction for both feeding and respiration is widely recognized (Hutchinson, 1973; Muller et al., 1982); however, dorsoventral compression and elongation of the lepisosteid skull relegates the opercle and subopercle to a shortened and narrow posterior margin relative to the length and width of the skull (Fig. 6). In *Atractosteus spatula*, opercular abduction alone is only capable of expanding the resting width of the cranium by ∼10% (Figs. 2 and 3). In contrast, suspensorial abduction in *A. spatula* expands the cranium more than 45% through lateral abduction of the cheeks (Figs. 2 and 3, Table 1). Lateral suspensorial movements that expand the opercular cavity were observed not only in feeding (Fig. 2e-h), but respiration as well (personal observation). These observations support a primarily hydrodynamic role of suspensorial abduction in gars over alternative hypotheses restricted to feeding – e.g., swallowing larger prey items or marginally extending the reach of the lateral tooth row during jaw closing.

Finally, the loose joint between the cleithrum and supracleithrum is particularly important for transitioning between the jaw opening and hyoid depression roles of the sternohyoideus during feeding. While at rest in a protracted state, the origin of the sternohyoideus on the cleithrum remains hidden behind the gill flaps, maintaining the streamlined profile of the body during lateral-snapping. In this position, the primary component of sternohyoideus-generated forces would be posteriorly directed, with little torque component. However, pectoral girdle retraction depresses the origin of the sternohyoideus below the ventral plane of the body, drastically changing the line-of-action of the sternohyoideus by increasing the torque moment-arm on the interhyal-ceratohyal joint (Fig. 8). Although the joint between the cleithrum-supracleithrum is likely a plesiomorphic feature of actinopterygians (Muller, 1987), in a dorsoventrally compressed taxon it takes on a particularly important role of modulating the nature of sternohyoideus activity throughout the feeding strike.

### Modulation of the feeding system with plesiomorphic hyoid musculature

Modulation of the feeding apparatus in gars into an anterior-to-posterior sequence of expansion requires the input of auxiliary muscles not typically associated with jaw opening. Anterior-to-posterior expansion of the feeding system is characteristic of most gnathostomes (Lauder & Shaffer, 1993) and also found in *Lepisosteus spp.* (Lauder, 1980a; Porter & Motta, 2004) and *Atractosteus spatula* (this study). However, gars lack the specialized jaw opening musculature of other groups of osteichthyans thought to be necessary for decoupling jaw opening from hyoid depression (Lauder, 1982; Wainwright et al., 2015; Wilga et al., 2000). *In situ* retraction of the pectoral girdle and sternohyoideus causes the entire feeding apparatus to expand as a single unit (Fig. 4c, Supplementary Video 1), but live feeding experiments show there is variation in onset of hyoid depression relative to jaw opening (see results), suggesting some component of active muscular control for this expansion. Although EMG data is not currently available for *A. spatula*, those data are available for *Lepisosteus* (Lauder, 1980a), and other studies have documented extensive conservation of EMG patterns between closely related taxa (Wainwright, Sanford, Reilly, & Lauder, 1989). The following is a proposed model for muscular inputs to the feeding system that explains the distinct, modulated phases of expansion during feeding in *A. spatula* (Fig. 8).

Jaw opening in *Atractosteus spatula* is the product of hyoid retraction while ventral or lateral rotation of the hyoid is restricted. Posterior movements of the hyoid are due to the sternohyoideus, which retracts the interhyal-ceratohyal joint and transmits posteroventral forces to the retroarticular process by means of the mandibulohyoid ligament, as in *Lepisosteus* (Lauder, 1980a). The extensive hyoid constrictor sheath of *A. spatula* (Fig. 6) acts as a muscular sling that allows hyoid retraction but maintains hyoid elevation and suspensorial abduction (Figs. 4b and 8b). In *Lepisosteus*, hyoid constrictors (interhyoideus, hyohyoideus, and adductor operculi) show a burst of activity during the earliest phases of jaw opening (Lauder, 1980a), and, while it was assumed this functioned to seal the opercular flap and prevent the influx of water through the gills, prior to hyoid depression (Fig. 2c) there is likely negligible negative pressure within the pharyngeal cavity. Conversely, these muscles may provide a means of coupling the powerful axial musculature to jaw opening – i.e., during neurocranial elevation, the hyoid is elevated away from the pectoral girdle (Figs. 2b and 8b, Supplemental Video 2), but sternohyoideus contraction causes the hyoid to retract relative to the suspensorium. Similar mechanisms have been described in various teleost groups, where neurocranial elevation coupled with pectoral girdle retraction can result in jaw opening with suspensorial adduction (Muller, 1987) or hyoid retraction and suspensorial abduction in tongue-raking taxa (Konow & Sanford, 2008). Brief hyoid elevation was noted in *Lepisosteus oculatus* (Lauder & Norton, 1980), and *Atractosteus spatula* demonstrates how hyoid constrictors, in restricting hyoid rotation during neurocranial elevation and hyoid retraction, could contribute to forceful mandibular depression and flat-plate suction.

The transition to hyoid depression and pharyngeal expansion in *Atractosteus spatula* is closely associated with pectoral girdle rotation due to powerful hypaxial musculature (Figs. 2d, 3b, 7b, and 8c, Supplementary Video 2). As a result of neurocranial elevation and pectoral girdle rotation, the origin and line-of-action of the sternohyoideus drop significantly below the initial center-of-rotation of the interhyal-ceratohyal joint (Fig. 8c). With relaxation of the hyoid constrictors and rotation of the pectoral girdle (Fig. 2d), force components of the sternohyoideus acting on the interhyal-ceratohyal joint necessarily shift as well (Aerts, 1991; van Leeuwen & Muller, 1984), decreasing the posterior force component and increasing torque. This helps to explain the characteristic dual firing pattern of the sternohyoideus recorded in *Lepisosteus*, in which the sternohyoideus is active during both jaw opening and closing, unlike *Amia* or *Polypterus* (Lauder, 1980a). However, muscle forces acting on the hyoid are not due to the sternohyoideus alone, and cleithral rotation enables hypaxial muscle forces to be transmitted to the hyoid by means of the sternohyoideus as well. Pectoral girdle rotation is an important mechanism of expansion of the feeding apparatus in *A. spatula* (Fig. 4d+h, Supplementary Video 1) and this study joins a growing body of literature documenting the contribution of hypaxial musculature to the generation of suction in actinopterygians (Camp, Roberts, & Brainerd, 2015; Muller, 1987; van Leeuwen & Muller, 1984).

Opercular expansion in *Atractosteus spatula* is greatly assisted by suspensorial abduction, which is the product not only of lateral rotation of the hyoid bars but also jaw closing. Suspensorial abduction occurs due to pectoral girdle rotation in *in situ* experiments, even without the input of the levator arcus palatini (Fig. 4g-h, Supplementary Video 1). The sternohyoideus is capable of transmitting the forces of pectoral girdle rotation to the hyoid, which posteroventrally depresses and laterally abducts the hyoid bars in a manner similar to models described by (Aerts, 1991; Muller, 1989; van Leeuwen & Muller, 1984), but in *A. spatula* peak suspensorial abduction coincides with jaw closure rather than peak gape. While the anterior hyoid bars are posteroventrally displaced by pectoral girdle rotation and the sternohyoideus, jaw closing causes the posterior ceratohyals to be *protracted* by means of the mandibulohyoid ligament (Supplementary Video 1). Maintenance of a fixed length mandibulohyoid ligament is recognized as playing a role in the unexpected movements of the lower jaws during jaw closing in sturgeons (Carroll & Wainwright, 2003) as well. This is perhaps the first study to propose a link between a direct contribution of jaw adductors to suspensorial abduction, potentially powering posterior expansion of the feeding apparatus during jaw closing.

### Evolution of osteichthyan feeding systems

The conclusion that hyoid constrictor musculature has an important secondary role in jaw opening in *Atractosteus spatula* helps to explain how specialized jaw opening muscles repeatedly evolved in multiple osteichthyan groups from the muscles that control hyobranchial and opercular motion. The main jaw opening muscle in *Amia* and teleosts is the levator operculi (Lauder, 1980a), which is a derived subdivision of the adductor operculi muscle (Lauder, 1982) and, according to current phylogenies (Grande, 2010), arose independently in *Amia* and teleosts. An independently derived levator operculi muscle with the potential to aid in jaw opening is also found in *Latimeria* (Lauder, 1980b). Similarly, the specialized depressor mandibulae muscles of lepidosiren lungfish and tetrapods appear to be convergently evolved from the dorsal hyoid constrictor sheet in those groups (Bemis, 1987). The repeated convergent evolution of specialized jaw muscles from muscles of hyobranchial and opercular control suggests a primitive, secondary jaw opening role of these muscles among all osteichthyans, allowing selective pressures to act on increasingly specialized jaw opening mechanisms in disparate groups. *A. spatula* demonstrates how hyoid constrictors may have functioned in this secondary capacity among basal osteichthyans prior to the evolution of specialized jaw opening subdivisions.

The synergistic interplay of jaw-ram and suction used by *Atractosteus spatula* helps to explain patterns of diversification among the closest extinct relatives of modern gars. While *A. spatula* likely utilizes the most suction among an otherwise heavily jaw-ram adapted clade, among extinct lepistosteids there are examples of extremely blunt-snouted forms for which suction-generation may have formed a greater component of their prey capture strategies (Brito, Alvarado-Ortega, & Meunier, 2017; Grande, 2010). With varied tooth morphology – durophagous (*Masillosteus*), “micro” (*Cuneatus*), and piscivorous (*Nhanulepisosteus*) – these taxa appear to have exploited various dietary niches and feeding strategies within the framework of the lepisosteid body-plan (Brito et al., 2017; Grande, 2010). As for the obaichthyid body-plan, *Dentilepisosteus* and *Obaichthys* possessed elongate rostra with short jaws (Grande, 2010), and appear to be convergently evolved with pivot-feeding syngnathiformes, which use an extreme form of jaw-ram followed by suction to capture prey (Longo et al., 2016). Elsewhere within the species-rich ginglymodian radiation, a diversity of body types and jaw morphology implies an even wider range of feeding strategies – durophagy, suction-feeding, non-predatory grazing, and detritophagy (Cavin, 2010; Grande, 2010; Schaeffer, 1967) – consistent with the findings of other researchers that show there is often widespread diversity of forms within clades favoring jaw-ram, suction, or a combination of both (Cooper et al., 2017; Longo et al., 2016). This diversity of fossil taxa, all more closely related to gars than other modern taxa (Grande, 2010), would not have been possible without an effective, plesiomorphic, suction-generating mechanism, maintained and demonstrated here by a “living fossil,” *A. spatula.*

## 5 CONCLUSIONS

Through the use of high-speed videography, contrast enhanced μCT, and *in situ* manipulations, this study documents a surprisingly versatile feeding mechanism of *Atractosteus spatula* that is capable of prey capture through a synergistic combination of jaw-ram and suction feeding. Examination of high-speed videos show *A. spatula* uses a combination of lateral jaw movements, flat-plate suction, and delayed hyoid depression to capture prey. Manipulation of fresh specimens shows that expansion of the feeding apparatus is accomplished by an elaborate linkage mechanism that joins together movements of the jaws, palate, hyoid arch, and pectoral girdle. Although cranial expansion was thought to be limited in gars, *A. spatula* demonstrates flexibility between elements of the suspensorium, hyoid arch, mandibular arch, and pectoral girdle that enhances expansive capabilities of this platyrostral taxon. Detailed anatomical reconstructions show that modulation of cranial expansion during the feeding strike is accomplished through the use of plesiomorphic hyoid constrictor musculature which decouples the dual functions of the sternohyoideus – jaw opening and hyoid depression – and demonstrates a likely primitive secondary role for muscles of hyobranchial and opercular control in assisting jaw opening in osteichthyans. Although gars are traditionally considered ram-feeders that only use suction for prey processing, these findings show lepisosteid feeding kinematics and morphology have been heavily influenced by the need to maintain suction while elaborating a lateral-snapping feeding mechanism.

## ACKNOWLEDGEMENTS

This study was made possible with specimens and assistance generously provided by the U.S. Fish and Wildlife Service, particularly William Bouthillier, Carlos Echevarria, and Haile Macurdy at the Warm Springs National Fish Hatchery, Georgia, where the young *Atractosteus spatula* specimens were raised. Thank you to the collection management staff at the Field Museum’s Division of Fishes who provided access to gar materials: Kevin Swagel, and Susan Mochel. Undergraduate assistants at the University of Chicago were instrumental in the completion of this project: Nora Loughlin and Leah Kessler assisted in collecting and analyzing alligator gar feeding sequences and aided with the general care of the animals.

---

**Supplementary Figure 1.**
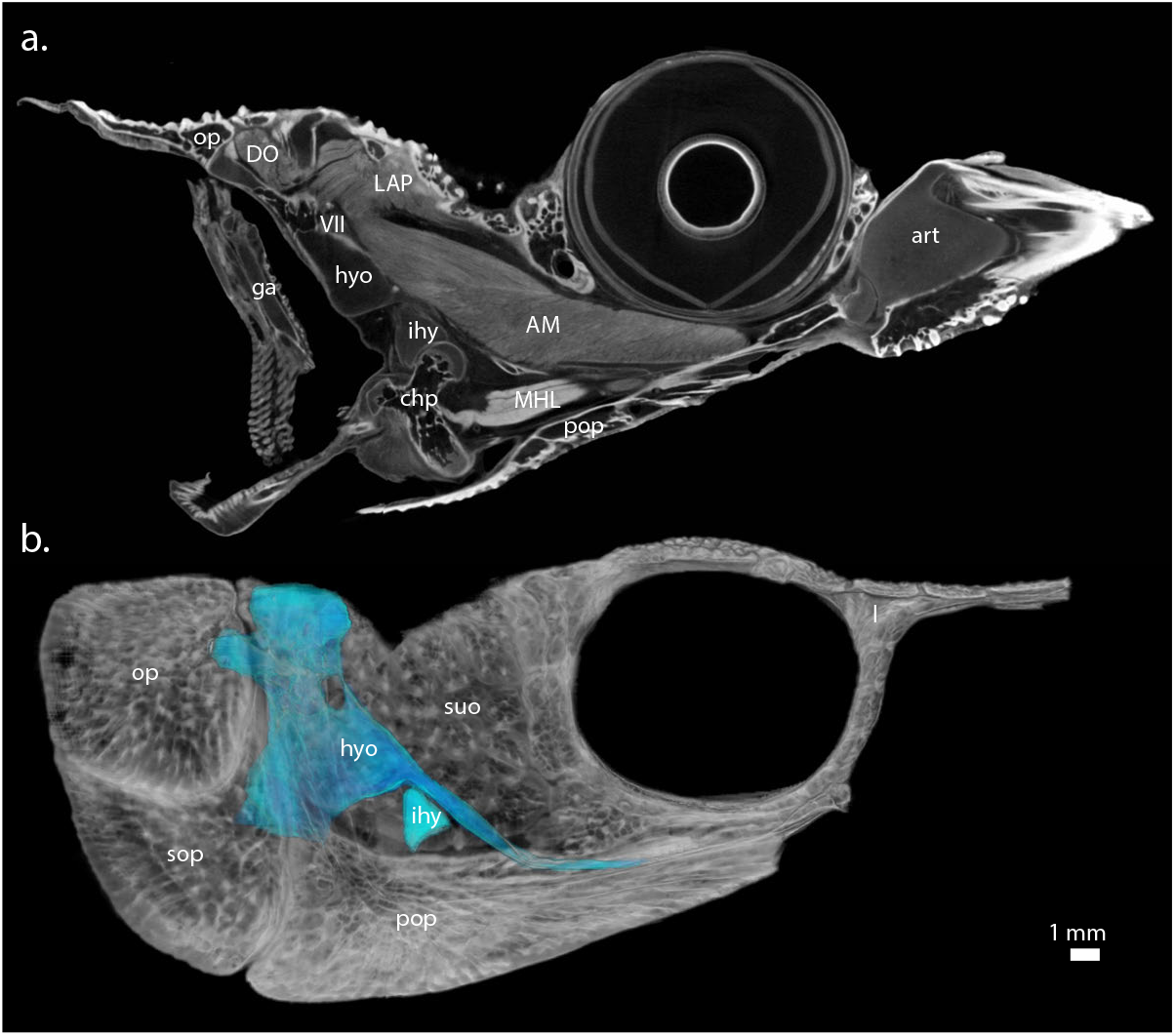
Morphology of the joint capsule containing the interhyal. Shown is the left joint capsule of the interhyal in (*a*) parasagittal cross section, and (*b*) medial view. This joint appears to be synovial, is bounded by the hyosymplectic (hyo) and preopercle (pop), and includes the posterior ceratohyal (chp), mandibulohyoid ligament (MHL), and branchiostegal rays. For abbreviations, see Table 2.

**Supplementary Figure 2.**
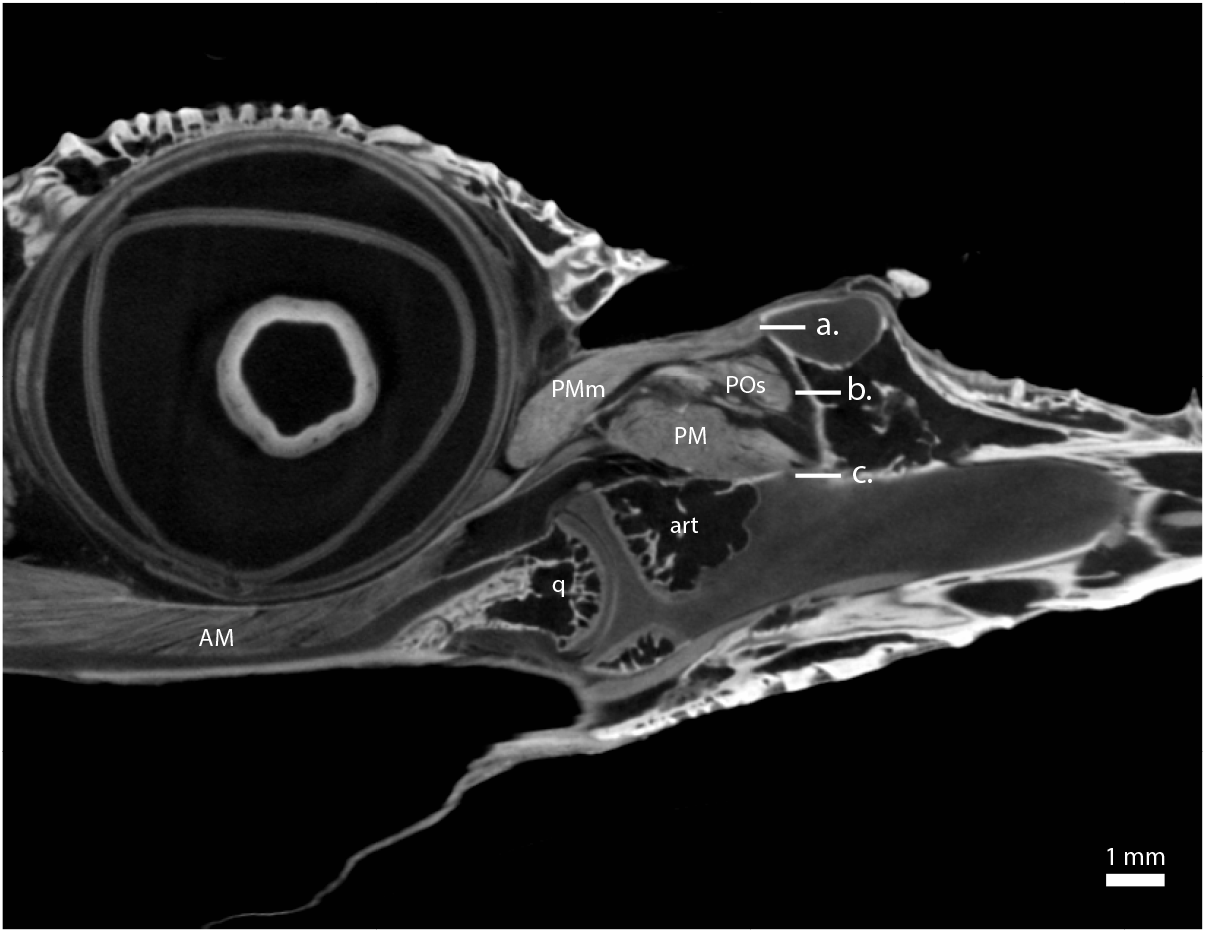
Adductor chamber of *Atractosteus spatula* showing multiple attachment areas for musculature. Jaw adducting musculature entering the adductor chamber attach to one of three areas of attachment: (*a*) dorsal, (*b*) middle, and (*c*) ventral. For abbreviations, see Table 2.

**Supplementary Figure 3.**
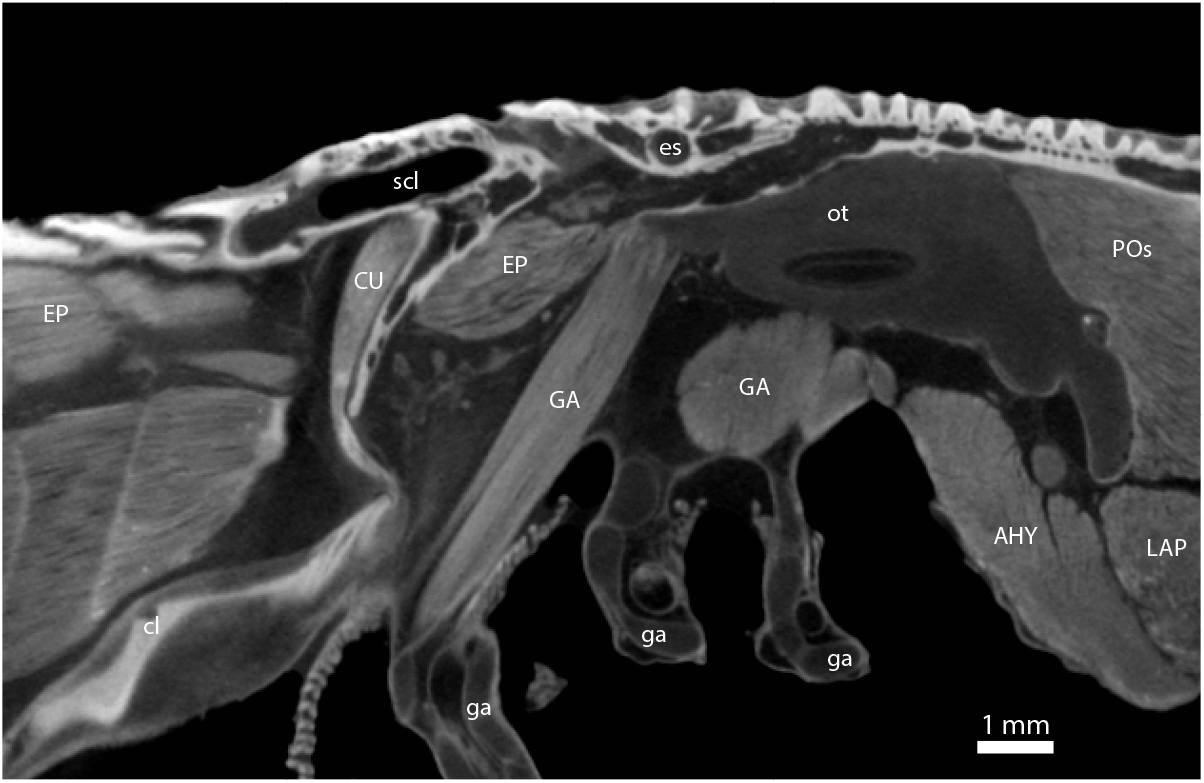
Parasagittal cross section of the head-shoulder interface. The supracleithrum (scl) is attached to the back of the skull (es); however, the cleithrum (cl) is only loosely connected to the supracleithrum by a flexible joint. A small slip of the cucullaris muscle (CU) originates on the medial surface of the supracleithrum and inserts onto the dorsoanterior prong of the cleithrum. For abbreviations, see Table 2.

**Supplementary Figure 4.**
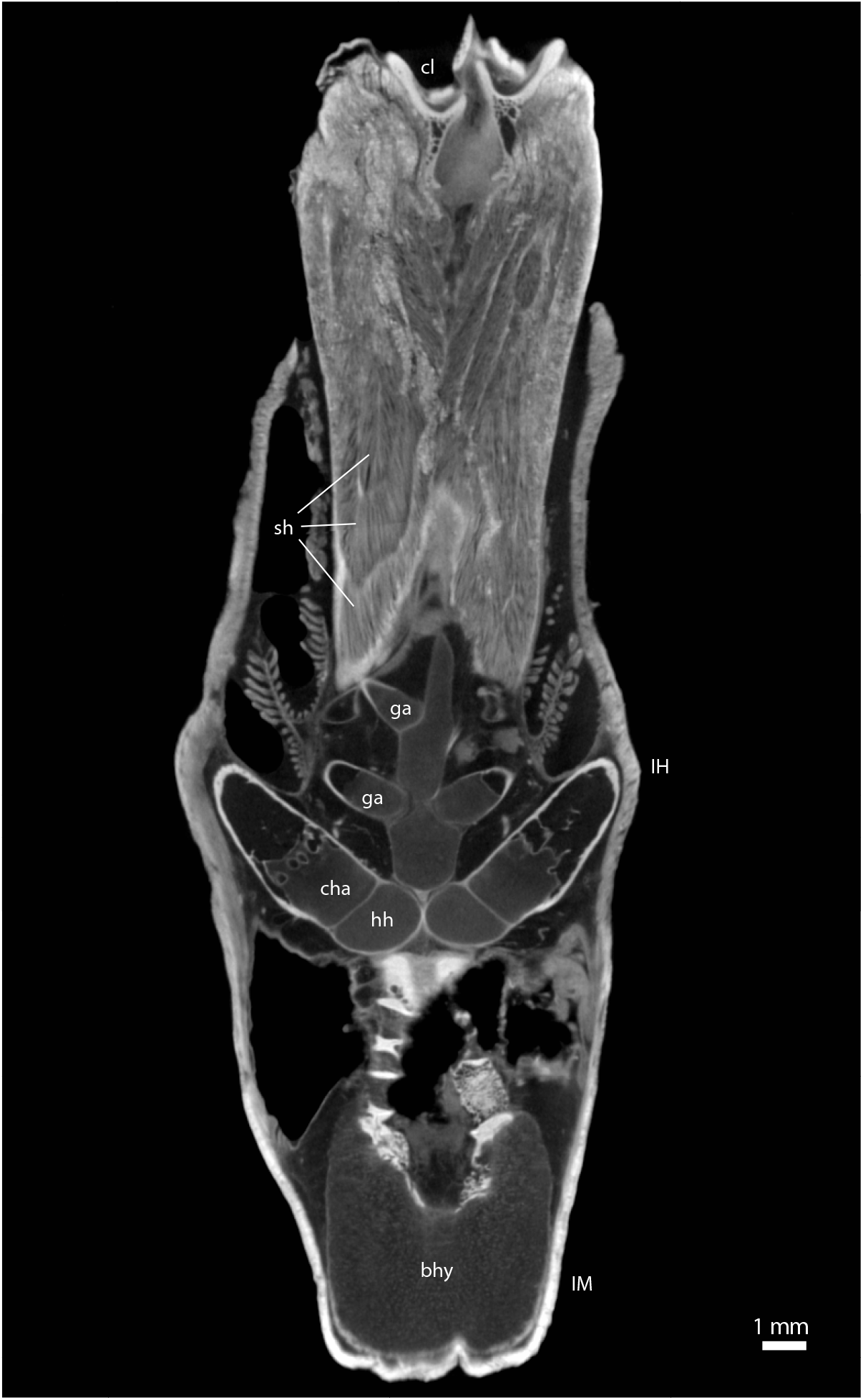
Divisions of the sternohyoideus muscle. Shown is a coronal section of the sternohyoideus muscle (sh), which is separated into at least three deeply nested cones along its length by septa internal to the muscle. Gill arches (ga) and the anterior ceratohyal (cha) lie dorsal to this muscle, which inserts onto the ventral surface of the hypohyals (hh) and basihyal (bhy). For abbreviations, see Table 2.

**Supplementary Figure 5.**
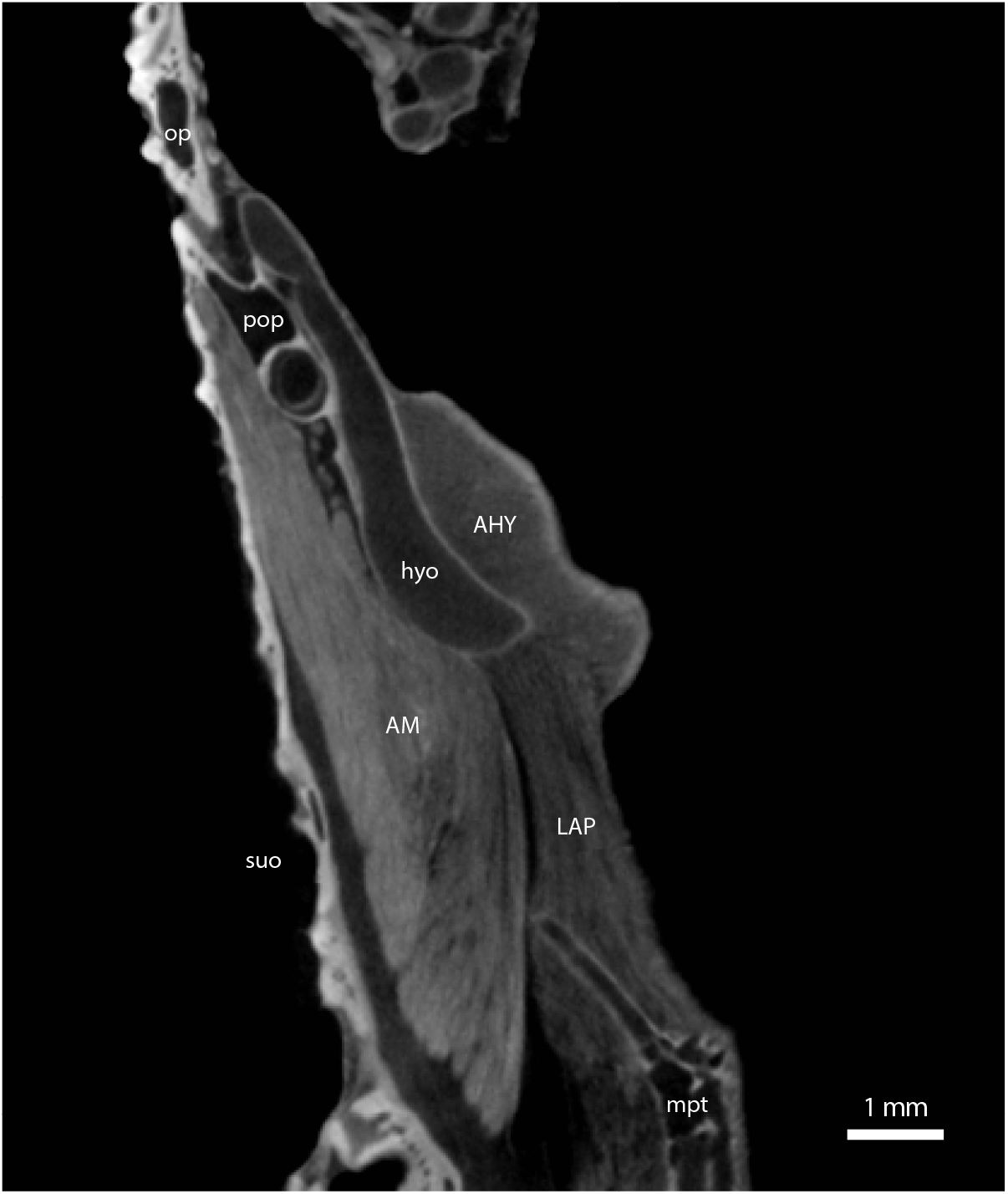
Small subdivision of the levator arcus palatini between the metapterygoid and hyosymplectic. The levator arcus palatini (LAP) originates on the sphenotic and primarily inserts onto the metapterygoid (mpt) with some fibers inserting onto the hyosymplectic. However, a small slip of this muscle directly connects the metapterygoid and hyosymplectic. Some muscle fibers of the adductor mandibulae (AM) also appears to originate from the hyosymplectic. For abbreviations, see Table 2.

